# Dynamic Gene-Based Ecophysiological Models to Predict Phenotype from Genotype and Environment Data

**DOI:** 10.1101/2021.02.07.429927

**Authors:** C. Eduardo Vallejos, James W. Jones, Mehul S. Bhakta, Salvador A. Gezan, Melanie J. Correll

## Abstract

Predicting the phenotype from the genotype is one of the major contemporary challenges in biology. This challenge is greater in plants because their development occurs mostly post-embryonically under diurnal and seasonal environmental fluctuations. Current phenotype prediction models do not adequately capture all of these fluctuations or effectively use genotype information. Instead, we have developed a dynamic modular approach that captures the genotype, environment, and Genotype-by-Environment effects to express the time-to-flowering phenotype in real time in *Phaseolus vulgaris*. The module we describe can be applied to different plant processes and can gradually replace processes in existing crop models. Our model can enable accelerated progress in diverse breeding programs, particularly with the prospects of climate change. Finally, a gene-based simulation model can assist policy decision makers in matters pertaining to prediction of food supplies.

## INTRODUCTION

Gregor Mendel (1865) deduced the genotype from the phenotype of garden peas. More recently, molecular characterization of Mendel’s genes (Reid and Ross 2011) has underscored the feasibility of using the genotype to predict the phenotype—the G2P connection. However, prediction of complex phenotypes remains among the biggest challenges in biology today, and particularly in plants because they develop post-embryonically (Steeves and Sussex 1989) under fluctuating environments resulting in different degrees of phenotypic plasticity (Scheiner *et al*. 2015). Crop simulation models (Jones *et al*. 2017; Wallach *et al*. 2018) and quantitative trait locus (QTL) analysis (Zhu *et al*. 2008; Bhat *et al*. 2016; Varshney *et al*. 2016) represent two complementary G2P approaches. While the former lacks a proper genetic framework, the latter doesn’t incorporate into the analysis the dynamic nature of the environment throughout the plant’s life cycle, despite existing dynamic methodologies (Wu and Lin 2006). These approaches have been combined previously with varying degrees of success (White and Hoogenboom 1996; Yin *et al*. 2000; Jones *et al*. 2003; Messina *et al*. 2006; Hammer *et al*. 2010; Bogard *et al*. 2014; Technow *et al*. 2014).

The realization that model parameter values are genotype-specific led to the idea that they contain genetic information (White and Hoogenboom, 1996), which could be extracted from the parameters to turn them into mathematical functions of underlying genetic factors. Parameter estimation and optimization procedures produce values that confer high model predictability, but the resulting model may not entirely reflect the genetic architecture of the trait. For instance, it has been shown that different parameter sets within a chosen model structure may be nearly as good as a selected set of parameters in reproducing the observed behavior of a system (Beven and Freer 2001; Acharya *et al*. 2017; Wallach *et al*. 2018).

However, model parameterization is based on some assumptions that may not be consistent with the genetic architecture of the associated biological process. For example, several models including APSIM (Hammer *et al*. 2010) and CROPGRO (Jones *et al*. 2003) estimate developmental rates based primarily on thermal time as modified by photoperiod requirements, and vernalization in some cases. These parameters are based on daily mean temperatures and cardinal temperatures. The assumption that all genotypes of a species have the same cardinal temperatures contrasts with reports of genetic variation for cardinal temperatures (Hardegree 2006; Rouan *et al*. 2018; Djanaguiraman *et al*. 2019). Also, computation of mean daily temperatures overlooks the effect of daily thermocycling on biological processes that affect yield (Peng *et al*. 2004; Lobell and Ortiz-Monasterio 2007; Shi *et al*. 2016). These observations suggest further testing of the hypothesis about the genetic information contained in model parameters.

Unlike traditional crop simulation models, statistical mixed-effects models offer an opportunity to predict phenotypic spectra using genotype (G), environmental (E), and GxE interactions data (Malosetti *et al*. 2013). For example, Bhakta *et al*. (2017) used only observational data to measure environmental and genetic effects to construct a static mixed-effects model for TTF in the common bean. These effects were estimated in the absence of assumptions about the functional forms and parameters for the TTF trait. This model used 12 QTLs along with average maximum and minimum daily temperatures (Tmax and Tmin), average day length (Day), average daily solar radiation (SRAD), one QTLxQTL and seven specific QTLxE interactions.

The high predictability of this model (Figure 1b and Table 1) indicated that statistical models represent a data-driven modeling approach that provides a robust foundation for the development of gene-based crop simulation models. The estimated genetic effects could be divided into three categories. The first were environmentally stable genetic effects; the second included the effects of specific QTLs on specific environmental responses, or GxE interactions, and the third comprised the effect of one gene on the action of another gene on the phenotype, or epistatic interactions.

**Table 1.** Comparative analysis of the performances of Cropgro-Bean, and the Static and Dynamic Mixed-Effects models.

| **Model** | **Count** | **MEff.** | **R<sup>2</sup>** | **R<sup>2</sup><sub>Adjusted</sub>** | **R<sup>2</sup><sub>Adjusted</sub>/R<sup>2</sup>** |
| --- | --- | --- | --- | --- | --- |
| Cropgro-Bean<sup>a</sup> | 123 | 0.955 | 0.961 | 0.843 | 0.874 |
| Std | 123 | 0.038 | 0.032 | 0.129 | 0.110 |
| Min | 123 | 0.745 | 0.845 | 0.379 | 0.449 |
| Max | 123 | 0.998 | 1.000 | 0.999 | 0.999 |
| Static M-E | 815 | 0.866 | 0.867 | 0.863 | 0.995 |
| Dynamic M-E | 815 | 0.911 | 0.912 | 0.909 | 0.997 |

<sup>a</sup>Mean values of performance for Cropgro-Bean; calculations were carried out independently for each RIL that had data at all five MET sites.

**Figure 1.**
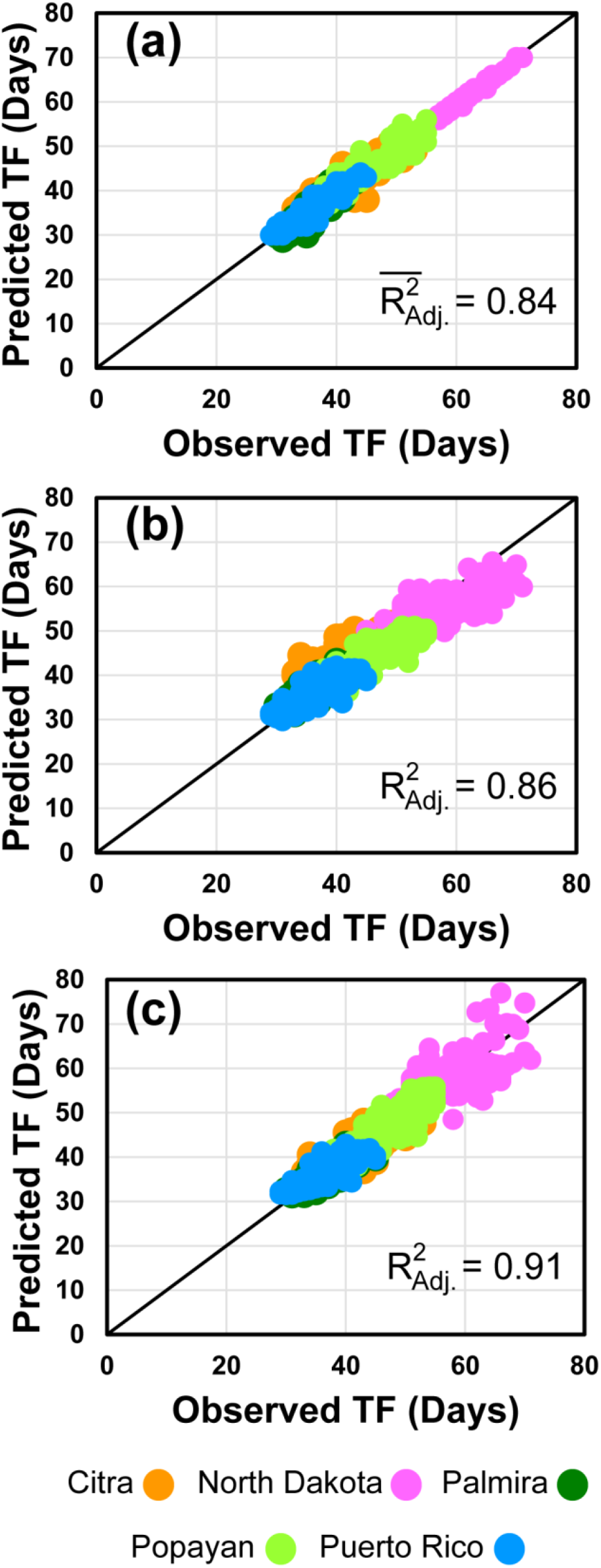
Observed vs. predicted time-to-flowering plots (1:1) of the experimental recombinant inbred family at the five sites. Predicted times were obtained with the CROPGRO-Bean model (**a**), the static (**b**) and the dynamic (**c**) statistical mixed-effects models.

The mesothermic range of environments of the experimental sites allowed Bhakta et al. (2017) to use linear functions for E effects. However, a more comprehensive effort will require a thorough survey of genotypic diversity through GWAS (Korte and Farlow, 2013), and an analysis of the widest possible range of environments, which will likely demand the use of nonlinear functions (Archontoulis and Miguez 2015). This approach will also require the converging efforts of geneticists, statisticians, physiologists, and process-based dynamic crop modelers. It must be pointed out that the mixed-effects model approach is completely different from phenotypic prediction methodologies based on the genomic selection method developed by Meuwissen *et al*. (2001), which uses all polymorphic markers for prediction purposes. We describe in this manuscript the conversion of a previously developed static mixed-effects model (Bhakta *et al*. 2017) into a dynamic model to predict the time-to-flowering (TTF) phenotype in *Phaseolus vulgaris* under varying environmental conditions.

## MATERIALS AND METHODS

### Plant material

The TTF phenotypes were collected from a RI family (n=188, F_11:14_) that was obtained from a bi-parental cross between a Mesoamerican and an Andean bean cultivar (Bhakta *et al*. 2017). Jamapa is a small black seeded bean cultivar from Mexico with an indeterminate growth habit and insensitive to photoperiod, whereas Calima is a large seeded and mottled Colombian bean cultivar with a determinate growth habit and sensitivity to photoperiod. The linkage map derived from this population was described previously (Bhakta *et al*. 2015) and the genotype data for the population can be found on line at https://figshare.com/s/50d1ddcaf8c04026dd4c.

### Experimental sites

Five geographical locations were selected to provide contrasting temperature and photoperiod conditions (Table S1). These included Citra, Florida (CIT); Prosper, North Dakota (ND); and Isabela, Puerto Rico (PR) in the United States; the Colombian sites Palmira, (PAL) and Popayan (POP) are near the equator provided short days, and their altitudinal difference (800 m) a temperature differential. Daily weather data from these sites are available in the Supplementary File *Weather_daily*.*txt*.

### Parameter estimation of genotype specific parameters (GSPs) for predicting anthesis day after planting (ADAP)

GSPs were estimated to predict ADAPs in the DSSAT CROPGRO-Dry Bean model (v. 4.5). These were planting to emergence (PLEM, thermal days), emergence to flowering (EMFL, photothermal days), and the slope of the relative response of development to photoperiod with time (photoperiod sensitivity, PPSEN, h^-1^). These GSPs were estimated for each genotype (parental and RILs) using a stepwise Markov chain Monte Carlo estimation approach as described previously (Acharya *et al*. 2017), but with minor changes. Briefly, PLEM was estimated first for each genotype across all five sites using emergence day after planting (EDAP) as the target output. For PLEM, the minimum and maximum values of the normal distributions for generating the proposal distributions were set to 2 and 20 thermal days, respectively. Next, EMFL was estimated using ADAP as the target output for each genotype across four of the five sites (i.e., all sites excluding ND) and the minimum and maximum distribution values were set to 20 and 40 photothermal days, respectively. Finally, PPSEN was estimated for each genotype with ADAP in ND as the target output (long-day site) with the minimum and maximum distribution values set to 0.001 and 0.500 h^-1^, respectively. Based on trace plots observed for all parameters, 3,000 iterations of GSP sampling (burn-in length set at 1,000 iterations) were sufficient for convergence. The critical short-day length (CSDL) below which reproductive development progresses with no day length effect (for short day plants), was set to the default value of 12.17 h.

### QTL mapping

QTLs that control GSPs associated with the TTF phenotype were mapped using composite (Zeng, 1993) and multiple (Kao *et al*. 1999) interval mapping as described elsewhere (Bhakta *et al*. 2017). The search for QTLs was performed with a window size of 5 cM, while a 50 cM distance was used as the minimum cofactor distance in the composite interval mapping scan. Threshold LOD values for QTL detection were determined at significance level of 0.05 using results obtained after 1,000 random permutations.

### Development of the mixed-effects and dynamic models

This model was derived from the one developed by Bhakta *et al*. (2017). However, instead of using TTF (days) as the observed response, we modeled the rate of development towards flowering (1/TTF (days)). The generic structure of the model based in *i* environmental variables and *j* QTL operators is:

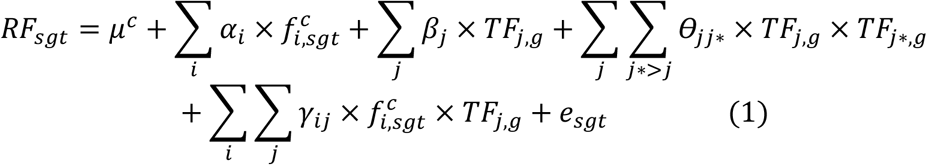

where *RF*_*sgt*_ (= *dP/dt*) is the rate of development for the genotype *g* on day *t* at site *s; μ*^*c*^ is the center, and *α*_*i*_, *β*_*j*_, *θ*_*jj\**_, and *γ*_*ij*_ are the model parameters to fit. In addition, the predictor variables are: 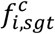 for the centered environmental factors, *TF*_*j,g*_ for the QTL operators, and their two-way interactions *TF*_*j,g*_ × *TF*_*j*,g*_ and *f*_*i,sgt*_ × *TF*_*j,g*_. Finally, *e*_sgt_ is the random residuals assumed to be a multivariate Normal distribution of the form ***e*** ∼ MVN(**0, R**), where **R** is a matrix of variance-covariance of unstructured form included to model the correlated nature of the observations belonging to the same genotype across sites. Note that the QTL operators for each genotype are coded as +1 for Calima alleles and –1 for Jamapa alleles. The above model was fitted using the strategy proposed by Malosetti *et al*. (2013), and further details were presented in Bhakta *et al*. (2017). The final model estimated the parameters for *i* = 4 environmental variables, *j* = 12 TF QTLs, a single G-by-G interaction and seven G-by-E interactions; the environmental variables represent the dynamic conditions under which the model operates. In a second step, the TTF of each RIL at each of the five experimental sites was predicted by first estimating the daily rates of development towards flowering (1/TTF (days)) by using as inputs the allelic makeup of the RILs at each of the 12 QTLs and the daily values of the four environmental variables (day length, solar radiation, and daily maximum and minimum temperatures). The TTF was identified when the daily integration of the daily rates reaches the value of 1.0.

A few general assumptions have been made for the dynamic mixed-effects model. The first is that responses to environmental effects are linear for any RIL within the range recorded during the MET for those variables (Tmax_sgt_, Tmin_sgt_, SRad_sgt_ and DL_sgt_). The model does not account for known nonlinearities in responses to these environmental variables that may occur in other environments. Likewise, it is assumed that there is a linear response to the genetic factors (12 QTLs) segregating in the bi-parental progeny.

The dynamic model also assumes that the daily developmental rate is controlled by the allelic makeup at the 12 QTLs of each genotype, and the daily weather conditions to which each genotype is exposed. This dynamic model fully applies to the bi-parental population for which it was constructed. It is also understood that the operational environmental domain of the model could be extended through the adoption of non-linear mixed-effect models, and that the versatility of such model within the crop species could be increased through the discovery of additional loci and alleles.

A two-part code for the statistical package R was written (Supplementary File) to carry out the procedures described above using the information contained in two data files (Supplemental data files). The ‘*Weather_daily*.*txt*’ file contains ten fields with the following headings: **SrNO**: Serial Number; **Site**: CIT=Citra, FL; ND=North Dakota, PAL=Palmira, Colombia; POP=Popayan, Colombia; PR=Puerto Rico; **Year:** The year in which the experiment was carried out – 2011 or 2012; **DOY**: Sequential day number starting with day 1 on January 01, which is also known as the Julian date; **DAP:** Number of calendar days after planting; **Srad:** Solar radiation in MJ m^-2^ d^-1^; **DayLhr:** Day length in hours; **Tmax:** Daily maximum temperature in degrees Celsius, °C; **Tavg:** Daily average temperature in degrees Celsius, °C; **Tmin:** Daily minimum temperature in degrees Celsius, °C.

The *‘R1data_weatherDAFtoFF*.*txt*’ file contains 19 fields with the following headings: **RIL:** The identifier of each RIL from the [Jamapa X Calima] cross; **Site:** Experimental site – CIT = Citra-FL; ND = North Dakota, PAL = Palmira-Colombia, POP = Popayan-Colombia; PR = Puerto Rico; **R1:** Number of calendar days when first anthesis was detected; **TF1 to TF12** (TTF QTLs in the RIL population); **Srad**_**m**_: Average solar radiation between planting and day of first anthesis for each RIL in MJ m^-2^ d^-1^; **DL**_**m**_: Average day length (h) between planting and day of first anthesis for each RIL; **Tmin**_**m**_: Average minimum temperature between planting and day of first anthesis for each RIL in degrees Celsius, °C; **Tmax**_**m**_: Average maximum temperature between planting and day of first anthesis for each RIL in degrees Celsius, °C. The TF QTLs and their chromosome and map positions (cM) according to the linkage map we constructed previously (Bhakta *et al*. 2015) are as follows: TF1, TF2, TF3 and TF4 (Chrom1, 22.1, 42.1, 58.8 and 70.0), TF5 and TF6 (Chrom3, 38.2 and 49.2), TF7 (Chrom4, 42.2), TF8 (Chrom6, 31.3), TF9 and TF10 (Chrom7, 11.7 and 98.7), TF11 and TF12 (Chrom11, 2.1 and 9.3). The numbers in the fields for TF1 to TF12 represent the operators for the Calima (+1) and Jamapa (−1) alleles.

In Part 1, the code estimates parameters for the mixed-effects model. Parts 2a and 2b constitute the dynamic components of the model. The Part 2a section estimates the daily rates of development towards flowering and an integrator adds up the daily progress until it reaches a value of greater than 1.02 for any given RIL at any given site. Part 2b calculates the day of flowering by interpolation using the integrated values flanking the value of 1.0. In addition to predicting TF for the experimental population, the code was included to do the same with a synthetic RIF that included all possible allelic combinations of the 12 TF QTLs. To work with the synthetic family, the first line of code for r1 must be turned into a comment and the following line must be activated by removing the comment symbol. After this change, the program will read the ‘*AllQTLcombo*.*txt*’ (Supplemental data file) instead of the ‘*R1data_weatherDAFtoFF*.*txt*’ file.

### Model Evaluation

Model efficiency (ME) is a measure of the predictive skill of the model. It expresses the model error on the basis of a predictor. At one extreme, if the prediction is a perfect one, then the efficiency is 100 %. However, if the average observation is used as the predictor and the there is no difference between the average and the predicted value, then the efficiency is zero.

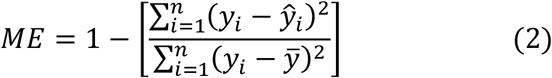

where *y*_*i*_ is the *i* ^th^observation, *ŷ*_*i*_ is the *i* ^th^ predicted value, and 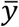 is the mean of the observed values.

The adjusted R^2^ is a modified value of the coefficient of determination that is adjusted for the number of predictors in the model and the size of the population. The R2adjusted decreases when the addition of parameters do not improve the fit of the model.

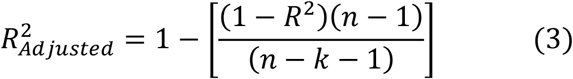

where R^2^ is the coefficient of determination, *n* is the number of samples and *k* is the number of parameters in the model.

### Development of gene-based time-to-flowering plug-in module

A computer program in FORTRAN was developed to produce a module for the dynamic simulation of the TTF, and to demonstrate the sensitivity of its prediction ability to both genetic and environmental variations. This dynamic module was designed to easily integrate into an existing ecophysiology model for beans, which simulates seasonal biomass growth, the timing of phenological events (including flower appearance), and final total and grain biomass (White *et al*. 1995; Jones *et al*. 2003).

This module has a structure that is similar to the dynamic module described above and integrates two sub-modules. The *Rate of Flowering Sub-Module* contains the daily rate of development Equation (Figure 2, Equation (5)), which computes both environmental and genetic input information. The *Driver Main Flowering Sub-Module* specifies the daily environmental variables and the genetic variables (QTL allele operators) for any particular genotype and passes them as inputs to the *Rate of Flowering Sub-Module* where the dynamic model is programmed. In turn, the daily rate and cumulative development, represented by *SumRF*_*i,j*_, is passed back to the driver main program. The module was designed this way so that it can be inserted into some other program, including a more comprehensive ecophysiology model like the CROPGRO-Bean model (Jones *et al*. 2003). The integrated operation of the sub-modules allow the simulation of TTF for different combinations of daily environmental conditions to create a type of sensitivity analysis to demonstrate some of the important variations that occur across environments for selected genotypes. For instance, the second sub-module could be made to read actual daily weather data and genetic variables for a simulation of one situation. Instead, this module specifies constant daily weather values, which are preselected to study how variations in any one variable (such as day length or temperature) influence rate of progress and time to first flower, a simple sensitivity analysis. This module also writes result files. The computer code and a brief summary of its modules (Rate of Flowering Sub-Module and Driver Main Flowering Sub-Module) can be found in the Supplementary File.

**Figure 2.**
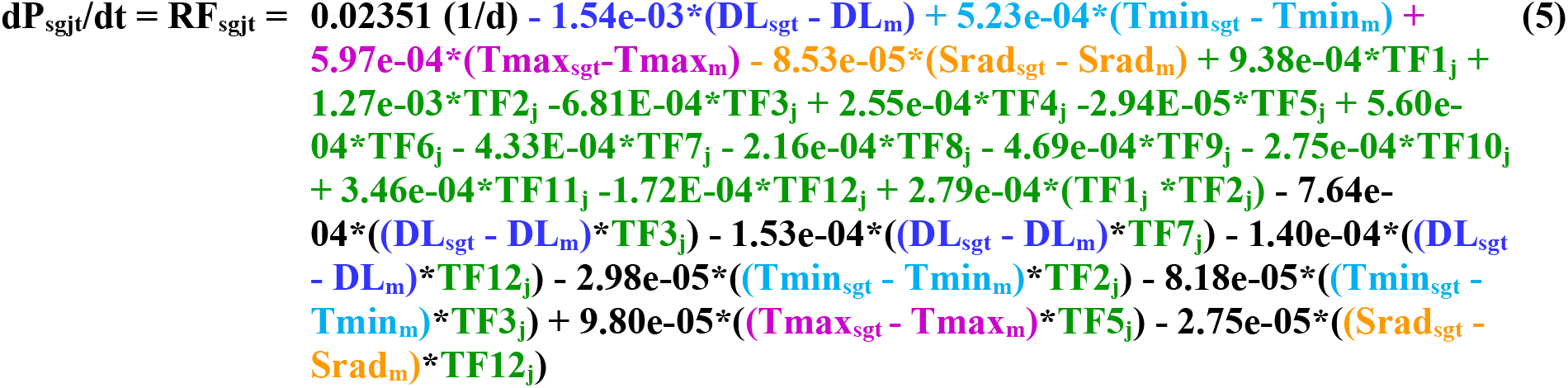
Differential equation describing the rate of development (RF) towards time-to-flowering as a function of genotype (g) and environmental (Tmax, Tmin, DL and Srad) factors at five experimental sites (s). This equation is represented by a linear mixed effects function where: DL_sgt_ = day length (hours) experienced by genotype *g*^*th*^ on day *t* at site *s*, DL_m_ = mean day length across all five sites (12.37 h); Srad_sgt_ = solar radiation (Srad, MJ m^-2^ d^-1^) experienced by genotype *g*^*th*^ on day *t* at site *s*, Srad_m_ = mean solar radiation across all five sites (18.218 MJ m^-2^ d^-1^); Tmin_sgt_ = minimum temperature (°C) experienced by genotype *g*^*th*^ on day *t* at site *s*, Tmin_m_ = mean minimum temperature across all five sites (16.128 °C); Tmax_sgt_ = maximum temperature (°C) experienced by genotype *g*^*th*^ on day *t* at ste *s*, Tmax_m_ = mean maximum temperature across all five sites (27.458 °C); TF1_j_ to TF12_j_ = QTL operators for the *j*^*th*^ allele (Calima alleles = +1 and Jamapa alleles = −1). For the rate equation: 0.02351 (d^-1^) is the mean effect of the environmental factors.

### Simulation Study Procedures

A simulation analysis was performed to explore variations among the lines as affected by the two most important environmental conditions in which the plants are grown. The gene-based time to flower model (**RF Module and Driver Module**) simulated the behavior of two RILs and the two parents of the RIJC population. Neither of the two parents were included in the bean MET dataset used to fit the model. The first RIL (RIJC031) was selected at random from the RILs in the population and the second RIL (RIJC366) was selected to have half of its QTL alleles different from the first one. These were arbitrarily selected to compare with the simulated responses of the two parents. Table S4 lists the QTL allele operators for these genotypes that were used to explore model responses to temperature and day length.

These four genotypes (Calima, Jamapa, RIJC031, and RIJC366) were simulated over a range of temperatures that represented the range that occurred across the five sites. For each simulation, day length and solar radiation were held constant so that only daily maximum and minimum temperatures were changed for each run but held constant for the duration of the simulation. The difference between daily maximum and minimum temperature was arbitrarily held constant at 8 °C. This resulted in a range of average daily temperatures that varied from 11 to 25 °C. For these simulations of temperature responses, day length was held constant at its mean value of 12.8631 h and solar radiation was held constant at its mean value of 18.2719 MJ m^-2^ d^-1^.

The four genotypes were then simulated 15 different times, varying day length for each one. Day length in each run was held constant whereas day length was varied among runs (between 11.5 and 18.5 h) (see the **Driver Module**). For the results presented for variations in day length, daily maximum and minimum temperatures were held constant at 26 °C and 18 °C, respectively. And, daily solar radiation was held constant for all of these runs as well at its mean value from the multiple environment trials (18.2719 MJ m^-2^ d^-1^). Results presented below were all based on simulations of the model. However, since the model described 91% of the variability in the observed data, they represent responses based on data, not on prior assumptions about response functions.

### Temperature sensitivity analysis

Simulated days to first flower varied considerably with temperature for each genotype (Figure S1). Results for the two parental genotypes (Calima and Jamapa) show similar responses to temperature, with time to flower for Jamapa being longer over the entire range of temperature variations. Differences between time to first flower at the 11 °C average temperature was about 20 days whereas it was less than 10 days at a mean temperature of 25 °C, indicating that Jamapa is more sensitive to cooler temperatures than Calima. The figure also shows that time to first flower for Calima dropped by about 32 days when mean temperature increased from 11 to 25 °C. In contrast, this decrease in time to first flower for Jamapa was about 47 days.

Figure S1 shows that one of the lines (RIJC031) was more sensitive to temperature variations than either of its parent genotypes. The response of RIJC366 to temperature was similar to that of Jamapa, but for each temperature, time to first flower was about 5 days shorter for RIJC366 than for Jamapa. The decrease in time to first flower for RIJC031 was about 69 days, about 50% more than either parent. This transgressive behavior was reported by Bhakta *et al*. (2017) and it is most likely due to the fact that one subset of Calima QTL alleles would lengthen the time to flower whereas the Jamapa alleles would have the opposite effect, and the second subset of QTLs has parental alleles with the opposite effects compared to the first subset.

### Day length sensitivity analysis

Both parents were sensitive to day length over the range of 11.5 to 15 h (Figure S2). However, time to flower under short days was shorter for Calima than for Jamapa (by about 6 days). Photoperiod exerted a significantly stronger effect in Calima than in Jamapa. Extending the day length from 11.5 to 18.5 h increased the time to flowering for Calima from 36 to 88 days, an increase of 52 days. In contrast, the same change in day length increased the time to flowering for Jamapa from 42 to 54 days, a 12-day delay showing that Calima sensitivity to day length was considerably higher than that of Jamapa. Responses of RIJC031 and RIJC366 were similar to that of Jamapa. However, time to flower of RIJC366 was shorter than that of RIJC031, which had a similar response to Jamapa.

### QTL effects on day length sensitivity

A final set of simulation runs was made to explore how changes in specific QTLs might affect responses to temperature and day length. We first identified QTLs that affected the response to day length based on the coefficients in the fitted model shown in Equation (5) (Figure 2). We selected TF1_g_, TF2_g_, and TF3_g_ for simulations. TF3_g_ has a direct effect on the response as day length is multiplied by the allele operator (Equation (5)); whereas TF1_g_ and TF2_g_ have an effect through multiple interactions.

The Jamapa response is controlled by all its QTL allele operator values (−1; Equation (5)), and was used as a baseline for comparisons. The first synthetic genotype, J-CTF3, was generated by replacing in Jamapa its TF3 allele with that of Calima (allele operator =+1; Table S4). Similarly, the second synthetic genotype (J-CTF1, 2, 3) was created by replacing in Jamapa its TF1, TF2, and TF3 alleles with the corresponding Calima alleles. Figure S3 shows that the response to day length of J-CTF3 exceeded that of Calima. In contrast, when the three QTLs alleles were replaced with Calima alleles the response to day length was similar to that of Calima. The discrepancy is most likely due to the multiple interactions involving TF2 and TF3.

## RESULTS

### Genetic Analysis of Crop Model Parameters

We selected the TTF trait to test the hypothesis that crop model parameters contain useful genetic information. We used the DSSAT CROPGRO-Bean model (Grimm *et al*. 1993) to model the TTF trait of beans using data from a multi-environment trial (MET) carried out at five sites (Fargo, ND; Citra, FL; Isabela, PR; and Palmira and Popayan, Colombia. Table S1) with a recombinant inbred (RI) family (F_11:14_, n = 188) of *P. vulgaris*, L. (Bhakta *et al*. 2015, 2017). The TTF trait was selected because it has high heritability (Buckler *et al*. 2009; Bhakta *et al*. 2017), and it is also very responsive to environmental variables (Greenup *et al*. 2009).

Like many crop models (Hadley *et al*. 1983; Yin *et al*. 1995; Jones *et al*. 2003; Zhang and Tao, 2013; Kumidini *et al*. 2014), CROPGRO (Grimm *et al*. 1993) handles daily environmental fluctuations using a development rate concept based on the following equation,

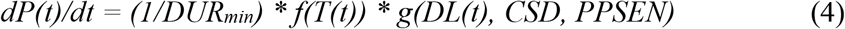

where *dP*(*t*)/*dt* is the daily rate of progress toward flowering at time *t* (in days), *DUR*_*min*_ is the genotype-specific parameter (GSP) representing TTF under optimal temperature (T) and day length (DL) conditions during the entire time period, *f(T(t))* is a T-dependent function that reduces developmental progress rate on day *t*, and *g* is a DL-dependent function that modifies the progress rate on day *t. CSD* and *PPSEN* are genotype-specific parameters (GSPs) that express genotype-specific sensitivity to DL; *CSD* is the genotype’s critical day length below which development is optimal, and *PPSEN* is the rate at which development rate is reduced for each hour above *CSD* (short-day plants). This rate is integrated over time (hours or days) until a developmental threshold is reached for simulating timing of a developmental transition such as TTF.

We applied inverse modeling (Acharya *et al*. 2017) to estimate the three GSPs associated with TTF (PL-EM: planting-to-emergence; EM-FL: emergence-to-flowering; and PPSEN: photoperiod sensitivity) for each RI line (Table S2); MET phenotypes and meteorological data can be found in the Supplementary Data File *Weather_daily*.*txt*. CROPGRO used these GSPs and daily environmental data from MET sites as inputs to simulate TTF. The observed vs. simulated plot (Figure 1a) appears to show that the model has high predictability, raising the possibility that flowering-associated GSPs may contain substantial genetic information.

QTL analyses of PL-EM, EM-FL and PPSEN in the RI family identified three, four and one QTL, respectively (Table S3). These QTLs didn’t explain all the observed variation, and only four of them co-localized with one third of QTLs detected through QTL analysis of the TTF phenotypes reported by Bhakta *et al*. (2017). This observation indicated that not all the information in GSPs is genetically tractable.

### Development of a Dynamic Mixed-Effects Model

As an alternative to the use of the traditional crop simulation model for predicting the phenotype from the genotype under varying environmental conditions, we revisited the mixed-effects model developed by Bhakta *et al*. (2017). A limitation of that model is its static nature because it uses the average values of E factors recorded during trait development. This approach cannot adequately represent the dynamic nonlinear and time-varying environmental effects on plant processes. However, the static model can be transformed into a dynamic gene-based process simulation module by applying the developmental rate concept described by Equation (4). An advantage of this application is that while TTF displays a curvilinear response to environmental factors, the rate of developmental progress exhibits a more linear response, at least under mesothermic conditions, resulting in an improved fit. This is basically the traditional crop modeling approach, which uses the duration of a developmental stage (TTF in days) to compute a rate of progress (RF(day^-1^) = 1/TTF) (Grimm *et al*. 1993; Yin *et al*. 2005). A fundamental assumption in this approach is that the rate of progress depends only on the environmental conditions of each day, and that these dependencies remain constant during the entire phase that is being modeled. Dynamic models are commonly divided into developmental/phenological phases with developmental transitions timed by this modeling approach.

We constructed a mixed-effects linear model of development rate (RF) as a function of G, E, and GxE interactions (See Equation (1) in the Methods section,). This approach models the trait as a developmental rate, rather than the occurrence of an event at a time point. Accordingly, the overriding assumption is that the genetic factors in the model are controlling a dynamic trait. Fitting the generic model to data from the MET yielded a function with 25 parameters (Table 2; Figure 2) describing the effects of 4 daily E variables, 12 QTLs, 1 QTLxQTL and 7 QTLxE interactions on the rate (Figure 1c). The relationship is centered on the mean E values recorded during the MET at the five sites. Dynamic implementation of the model requires daily (*t*) computation of RF_s,g,t_ using the daily values of each of the *i* E factors experienced by genotype *g* at each experimental site (*s*), and NOT the average values over the vegetative phase of each RIL at each experimental site as implemented in the model developed by Bhakta *et al*. (2017).

**Table 2.** Estimated model parameters from differential equation describing the rate of development (RF_sgt_) towards time-to-flowering as a function of genotypes QTL operators (TF_g,j_) and environmental factors (f_i,sgt_ = DL, Srad, Tmax and Tmin) at five experimental sites (s). Predictor variables are: DL_sgt_ = day length (hours) experienced by genotype *g*^*th*^ on day *t* at site *s;* Srad_sgt_ = solar radiation (Srad, MJ m^-2^ d^-1^) experienced by genotype *g*^*th*^ on day *t* at site *s*; Tmax_sgt_ = maximum temperature (°C) experienced by genotype *g*^*th*^ on day *t* at site *s*; Tmin_sgt_ = minimum temperature (°C) experienced by genotype *g*^*th*^ on day *t* at site *s*; TF1_g_ to TF12_g_ = QTL operators coded as +1 for Calima alleles and –1 for Jamapa alleles.

| Variable | Parameter | Estimate | Description |
| --- | --- | --- | --- |
| **Center** | $\mu^c$ | 0.0235198065831426 | Overall Center |
| **$DL_{sgt}$** | $\alpha_1$ | -0.0015423214758081 | Environmental Factors |
| **$Srad_{sgt}$** | $\alpha_2$ | -0.0000853054955707 | |
| **$Tmax_{sgt}$** | $\alpha_3$ | 0.0005966415621148 | |
| **$Tmin_{sgt}$** | $\alpha_4$ | 0.0005228804696115 | |
| **$TF1_g$** | $\beta_1$ | 0.0009382848209270 | QTL Operators |
| **$TF2_g$** | $\beta_2$ | 0.0012733168457595 | |
| **$TF3_g$** | $\beta_3$ | -0.0006810089804720 | |
| **$TF4_g$** | $\beta_4$ | 0.0002546737990060 | |
| **$TF5_g$** | $\beta_5$ | -0.0000294375056886 | |
| **$TF6_g$** | $\beta_6$ | 0.0005598758808943 | |
| **$TF7_g$** | $\beta_7$ | -0.0004332155371734 | |
| **$TF8_g$** | $\beta_8$ | -0.0002159196193359 | |
| **$TF9_g$** | $\beta_9$ | -0.0004693200551048 | |
| **$TF10_g$** | $\beta_{10}$ | -0.0002745608987544 | |
| **$TF11_g$** | $\beta_{11}$ | 0.0003463304080702 | |
| **$TF12_g$** | $\beta_{12}$ | -0.0001723147103969 | |
| **$TF1_g \times TF2_g$** | $\theta_{12}$ | 0.0002794695951983 | Interaction: QTL by QTL |
| **$DL_{sgt} \times TF3_g$** | $\gamma_{13}$ | -0.0007636661891134 | Interaction:<br>Environmental Factor by<br>QTL Operator |
| **$DL_{sgt} \times TF7_g$** | $\gamma_{17}$ | -0.0001527383245705 | |
| **$DL_{sgt} \times TF12_g$** | $\gamma_{112}$ | -0.0001400623472723 | |
| **$Tmin_{sgt} \times TF2_g$** | $\gamma_{42}$ | -0.0000298150506285 | |
| **$Tmin_{sgt} \times TF3_g$** | $\gamma_{43}$ | -0.0000818315988969 | |
| **$Tmax_{sgt} \times TF5_g$** | $\gamma_{35}$ | 0.0000980123505664 | |
| **$Srad_{sgt} \times TF12_g$** | $\gamma_{212}$ | -0.0000274938204046 | |

Thus, day *t* on which genotype *g* first flowers at site *s* is determined by integrating the rate of development function (Figure 2) over time until it computes a threshold value of 1.0 for *P*_*s,g,t*_ (unitless) according to:

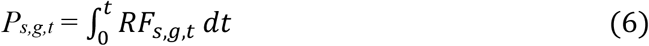

where *P*_*sgt*_ = 0 at *t* = 0 (planting day). The assumption of linearity of E effects on developmental rate is reasonable for environments that do not reach lower or upper limits where the responses are highly nonlinear. This is the main reason for centering the model on the E factors, and also to adequately represent the effect of the multiplicative terms (QTLxQTL and QTLxE) over the range of data in the MET. The dynamic model produced a highly accurate estimation of the TTF phenotype using genetic (QTL) and environmental (E) inputs.

### Comparative Analysis of Model Performance

We compared the performances of CROPGRO, the static mixed-effects model developed by Bhakta *et al*. (2017), and the dynamic model described here for their abilities to simulate the TTF phenotype in the common bean (See Table 1). For this purpose, we identified 123 RILs that had observations at all 5 MET sites. CROPGRO requires the estimation of three model parameters (GSPs) for each RIL to simulate TTF. For this reason, evaluations were carried out independently for each RIL. While crop model parameters had to be estimated for each genotype, mixed-effect model parameters were estimated for the entire population and consequently, data from a total of 815 RIL-site combinations were used for model evaluation.

Model efficiency (Equation (2)) estimates for CROPGRO averaged 0.955, with a range of 0.745 to 0.998, and a small standard deviation. This suggested that the variation not explained by the model was relatively small compared to the variation in the observed values. Model efficiencies in the static and dynamic models were lower than the CROPGRO average, 0.866 and 0.911, respectively, suggesting that CROPGRO is a better model, on average. However, the wide range of variation in the prediction of TTF by CROPGRO indicated a component of uncertainty and required further assessment.

To conduct a further evaluation of the models we used the R^2^_adjusted_ (Equation (3)) value because it makes adjustments for both population size and for the number of model parameters. Accordingly, CROPGRO produced a total of 369 parameter values resulting in 615 simulations. The R^2^_Adjusted_ values of the RILs averaged 0.843 with a range between 0.379 and 0.999. In contrast, the static and dynamic mixed-effects models used only 25 parameter values for the entire populations grown at the five MET sites. Thus, these parameters simulated 817 RIL-site combinations with R^2^_Adjusted_ values of 0.863 for the static and 0.909 for the dynamic models.

Factoring in sample size and the number of parameters showed that the average R^2^_Adjusted_ value for CROPGRO simulations represented a 0.87 fraction of the R^2^, with a range of 0.449 to 0.999. In contrast, the R^2^_Adjusted_ values of the static and dynamic mixed-effects models were a 0.99 fraction of their respective R^2^ values indicating that the number of parameters did not artificially increase model performance.

### Validation and Sensitivity Analysis

Validation of the dynamic model was carried out with two data sets. The first set comprised the parental genotypes that were grown at the five MET sites but were not included in the construction of the model (Figure 3a). The R^2^_Adjusted_ value was not calculated for this set because the number of samples was lower than the number of parameters. However, a Model Efficiency of 0.99 indicated that the variation between prediction and observation was a very small fraction of the variation among the observations. The second set included data from a planting of 100 of the RILs along with the parents in 2016 in Citra, FL (Figure 3b). Model prediction in the 2016 Citra trial (Figure 3b) showed low performance with an R^2^_Adjusted_ value of 0.52. These results are similar, to some extent, to those reported for North Dakota (part of the MET), and the causative factors are likely the slight high temperatures and longer days. However, the root mean squared error for the parental lines was of 1.5 days and that for the 2016 Citra experiment 2.4 days indicating the model had a reliable prediction ability.

**Figure 3.**
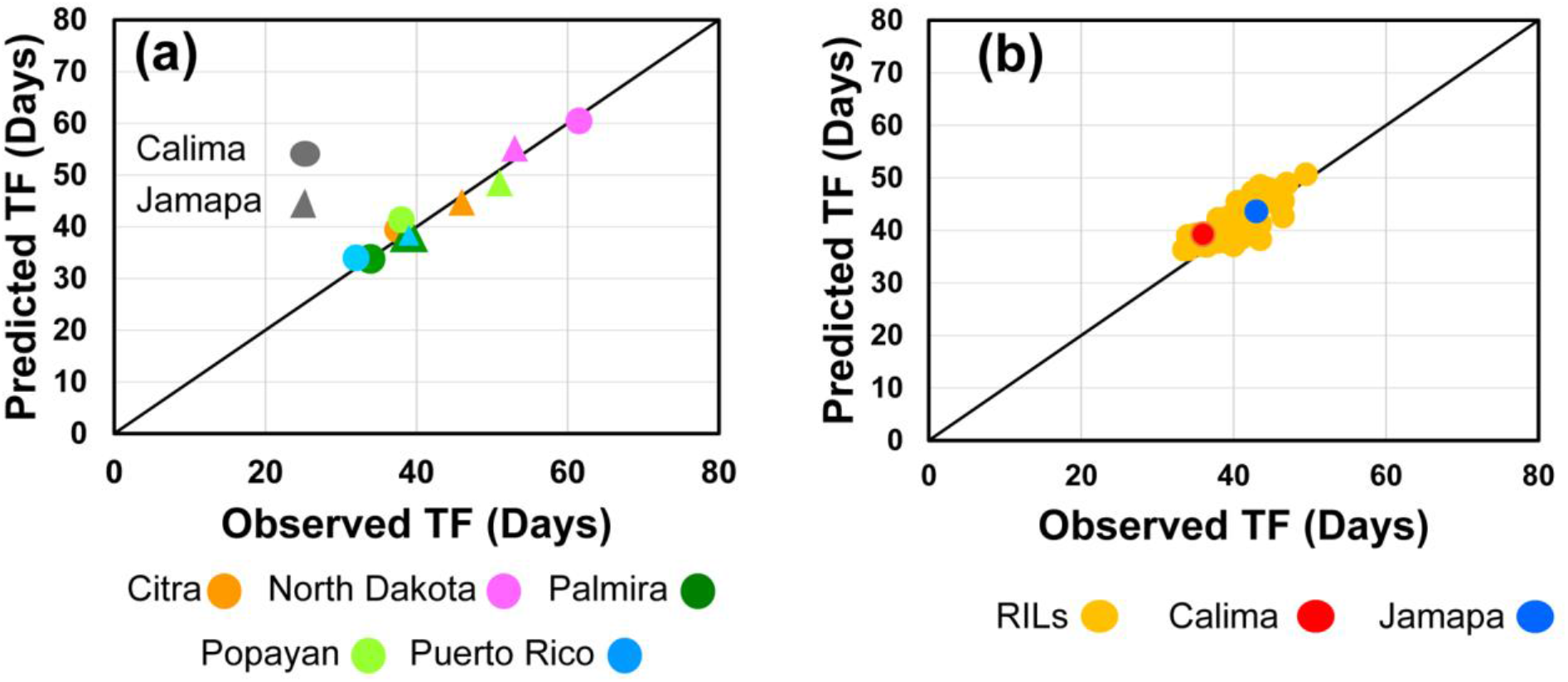
Validation tests of the RF dynamic simulation model. Observed vs. predicted plots (1:1) of two data sets not involved in generating of the model. (**a**) Parental genotypes that were grown along the RI family at the five MET sites (Model Efficiency = 0.997; RMSE = 1.54 days). (**b**) A subset of 100 RILs, along with the parents, grown in Citra, FL, in 2016 (Model Efficiency = 0.36; RMSE = 2.44 days; R^2^_Adjust._ = 0.52).

Sensitivity analysis of the flowering module analyzed the range of combinations of environmental variables (Tmax_*s,t*_, Tmin_*s,t*_, Srad_*s,t*_, and DL_*s,t*_) for different genotypes for each day *t* after planting. Environmental variable ranges were restricted to those observed in the MET, but with small extrapolations in the values to study the overall patterns of effects. The simulated effect of average daily temperature ((Tmin + Tmax)/2) on TTF for the two parental lines displays the typical curvilinear shape, which is due to the linear effect of temperature on developmental rate (Figure 4a). Dropping the average daily temperature from 25 °C to 10 °C increased Calima’s TTF from 34 to 72 days under short days (11.5 h), and from 40 to 111 days under longer days (13.5 h); no flowering occurred within 200 days at the lowest temperature for days longer than 15 h. In contrast, the temperature sensitivity of TTF in Jamapa was not as greatly affected by photoperiod as it was in Calima. Day length simulations, holding Tmin and Tmax constant at 18 and 26 °C, respectively, showed that long days significantly delay TTF in Calima as compared to Jamapa (Figure 4b), in agreement with previous observations (Bhakta *et al*. 2017).

**Figure 4.**
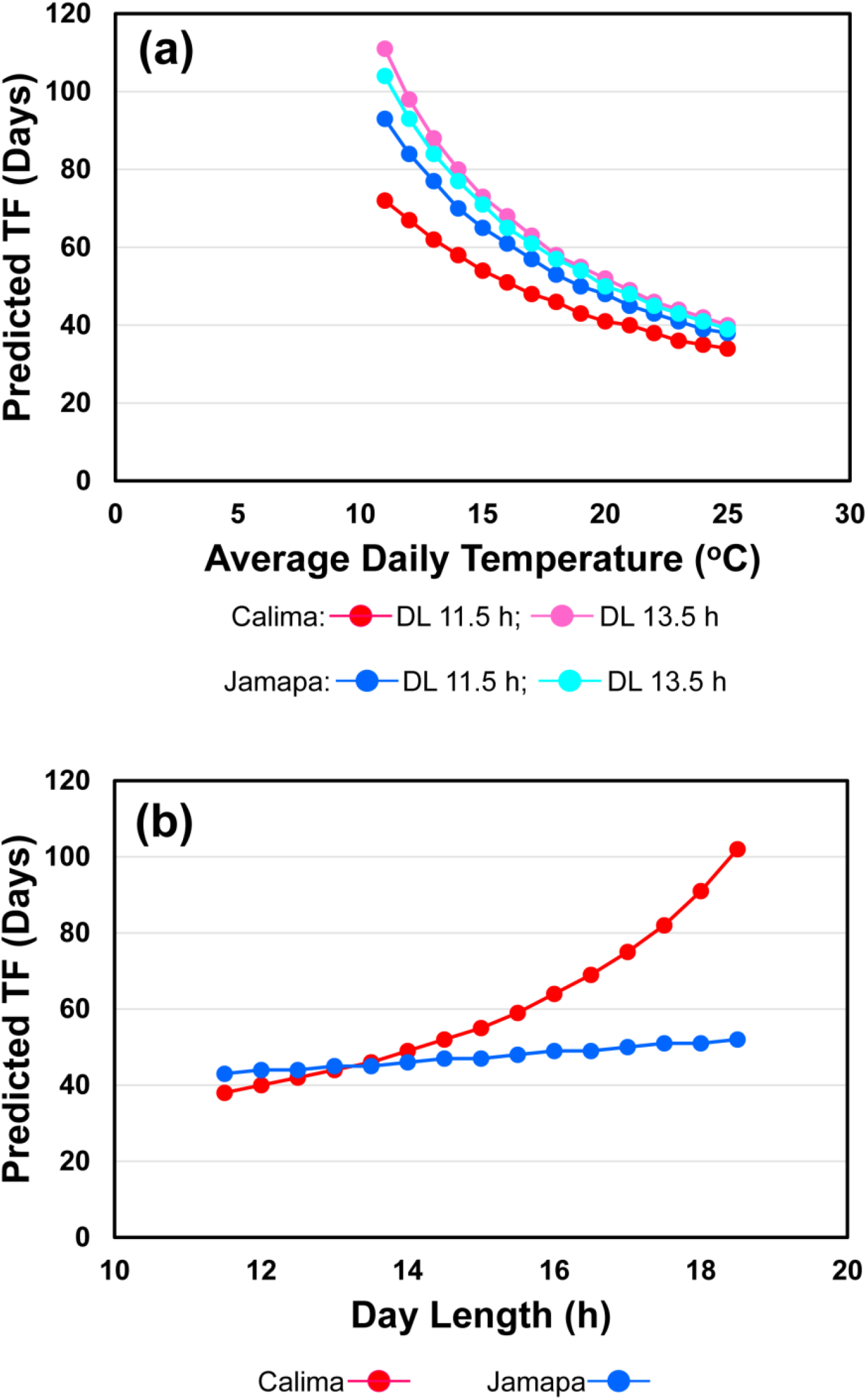
Sensitivity analysis of the Time-to-Flowering module using the parental genotypes: Calima and Jamapa. (**a**) Simulations of temperature effects on TF according to Equations (5) and (6) under two day lengths. (**b**) Simulation of day length effects on TF with Tmax and Tmin values of 26 and 18 °C, respectively.

## DISCUSSION

Testing the hypothesis that crop model parameters contain genetic information produced mixed results. On the one hand, we detected some QTLs for model parameters controlling the TTF phenotype in CROPGRO, but on the other hand those QTLs did not explain all the variation observed for this trait and did not completely match the all the QTLs detected by Bhakta et al. (2017). These results are very similar to those reported by Bogard *et al*. (2014), who detected high predictability (r^2^=0.97) of heading time by the wheat-APSIM model, which was reduced significantly (r^2^=0.68) when model parameters were turned into linear functions of QTLs. Taken together, these results indicated that not all the information in model parameters is genetically tractable. Crop simulation models were not developed within a genetics framework, but with an exclusive focus on the environmental dependencies of plant processes, while model parameters were devised to represent these dependencies.

The comparative analysis of the models showed that, on average, CROPGRO appears to have a high predictive ability. However, the variability of TTF prediction in the RI family, as shown by the wide range of R^2^_Adjusted_ values (0.379 - 0.999) showed a strong component of unpredictability. This is further underscored by the range in the R^2^_Adjusted_/ R^2^ ratios (0.45 – 0.99). These observations suggest that some model assumptions may not necessarily apply to all genotypes. For instance, most models including CROPGRO assume cardinal temperatures are fixed in a species. However, if the range of temperatures for biological activity of individual genotypes vary significantly, then overlooking this type of variation may create challenges for models that do not consider this phenomenon at all.

The results presented in this manuscript clearly indicate that statistical mixed-effects models can be used effectively to predict the phenotype using genotypic (G) and environmental (E) data inputs. Unlike other models, this model was developed without assuming that all genotypes have the same temperature response; furthermore, it included significant and specific GxE interactions. An important difference between the crop model-genetics approach and the mixed-effects approach is as follows. While the crop model approach used derived phenotypes, that is, estimated model parameters, to obtain QTL information, the mixed-effects model used QTL information obtained from genetic analysis of the primary phenotype. These results also demonstrated that the overall approach used in many crop models for computing rate of progress toward first flowering can be used in a statistical approach in which G, E, and GxE effects are considered. The increase in model accuracy of the dynamic model (R^2^_Adjusted_ = 0.909) over the static model of Bhakta et al. (2017) (R^2^_Adjusted_ = 0.863) can be ascribed to the use of daily E input values instead of average E factor values over a period of time.

Phenology modules play a key role in crop models because they set the rate of development for the crop, which can significantly affect productivity by altering the dynamics of assimilate partitioning throughout the crop’s life cycle. However, phenology modules are typically difficult to parameterize because they are commonly affected by GxE interactions. Thus, the inclusion of specific GxE interactions in the dynamic mixed-effect TTF model makes it a potential key module component of a comprehensive crop model. Results of the sensitivity analyses with the parental genotypes and few selected RI lines along with the validation exercises at the five experimental MET sites provide strong support for the TTF dynamic model, but the validation exercise obtained with the 2016 Citra data (Figure 3b) indicated that perhaps that effect high temperatures in combination with long days still need to be fully captured by the model.

The construction of a dynamic mixed-effect model that uses G and E data as sole inputs, as described above, could be applied to other plant processes in crop species in general. This raises the possibility of developing dynamic gene-based crop simulation models (DGCSM) capable of integrating information from interconnected phenology and growth process-oriented modules. Growth process modules trace resource acquisition of different organs. For example, a leaf area expansion module could integrate input from sub-modules for rate of leaf appearance, individual leaf area expansion, branch appearance, and similar branch sub-modules. This approach accentuates the increased granularity required by DGCSM, a characteristic that makes the phenotype more genetically tractable and reduces equifinality problems in phenotype prediction; for instance, two genotypes with the same leaf area – one with few large leaves and the other with many small leaves. Furthermore, determining the identities of QTLs can create a powerful connection between DGCSM and dynamic gene regulatory networks, and therefore establish a G2P bridge across scales of time and space.

Current crop models can be gradually converted into DGCSM by replacing existing process model components with modules that incorporate G, E, and GxE relationships. It represents an innovative approach that combines genetics and modeling to increase both prediction effectiveness and understanding of genetic effects on biological processes. We have integrated a stand-alone two-module dynamic simulation program into the DSSAT CROPGRO model (Oliveira *et al*. 2021; see also Supplementary File). The first module computes the daily rate of development according to the fitted function (Table 2; Figure 2) and requires the QTL allele operators of individual genotypes and daily environmental values (Tmax_*i*_, Tmin_*i*_, Srad_*i*_, and DL_*i*_). The second module integrates the daily rates over time according to Equation (6) to predict time to first flower. The code can be extended to a wider genotypic base and environmental range after proper estimation procedures.

To illustrate the usefulness of DGCSM in plant-breeding, we simulated the phenotypes of multiple QTL allelic combinations grown at each one of the five experimental site. These are all possible allelic combinations (2^12^ = 4,096) of the 12 QTL segregating in our experimental RI family (Figure 5). Such simulation enables evaluation and selection of suitable allelic combinations that produce a desirable TTF phenotype in specific environments, which may include those projected for climate change. DGCSM can expedite genetic progress by reducing the need for expensive MET. Furthermore, as an aid in precision breeding, DGCSM can be turned into expert systems to search the gene space (QTL database) to create suitable ideotypes (Donald, 1968; Rötter *et al*. 2015). Finally, projected increases in human population and climate change highlight the urgency with which worldwide food security needs to be addressed (Godfray and Garnett, 2014). DGSCM in conjunction with climate models could assist policy decision makers in developing information about food supply futures to improve worldwide stability.

**Figure 5.**
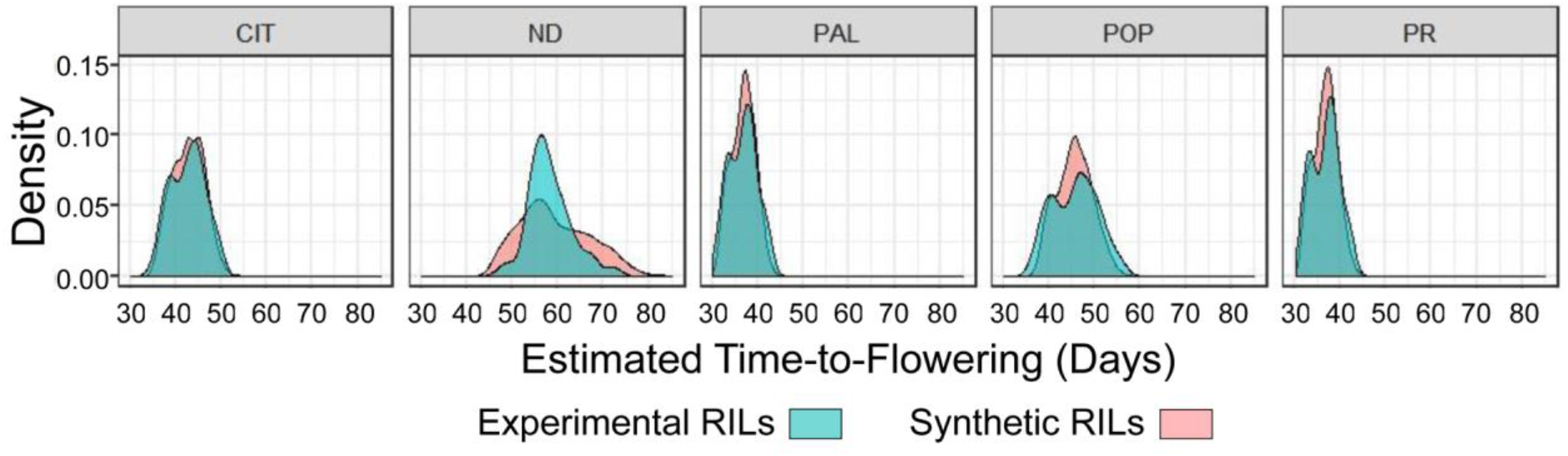
Density plots of Time-to-Flowering estimate of the RI families at the five experimental sites. Frequency distribution of the experimental RI family (Green; n = 188) and the synthetic RI family comprising all 4,096 possible allelic combinations of the 12 QTLs as simulated by the dynamic module. These plots indicate that multiple allelic combinations can attain the same phenotype providing a choice of convenience to plant breeding programs.

## SUPPORTING INFORMATION

The following additional information is available in the online version of this article.

**Table S1**. Geographical and environmental characteristics of the five experimental sites.

**Table S2**. Genotype-specific parameter (GSP) values.

**Table S3**. Summary of QTL mapping results.

**Table S4**. Allele operators of the TF QTLs.

**Figure S1**. Temperature simulations generated with the dynamic gene-based time-to-first-flower bean model – implementation of Eqs. (2) and (3).

**Figure S2**. Photoperiod simulations generated with the dynamic gene-based time-to-first-flower bean model – implementation of Eqs. (2) and (3).

**Figure S3**. Photoperiod simulations of real and synthetic genotypes created through substitutions of QTL alleles.

**Supplementary Text**

**Flowering Rate Model-R code**

**Rate of Flowering Sub-Module – Rate of Progress toward First Flower**.

**Driver Sub-Module**.

**Supplementary Data Files Weather_daily.csv**

**RIL_R1sata_weatherDAPtoFF.csv AllQTLcombo.csv**

## Supporting information

AllQTLcombo.csv

RIL_R1dataDAPtoFF.csv

Weather_daily.csv

## ACKNOWLEDGEMENTS

We acknowledge the invaluable contribution to data collection made by Juan Osorno and Raphael Colbert at North Dakota State University, James Beaver and Abiezer Gonzalez at the University of Puerto Rico, Idupulapati Rao, Stephen Beebe and Jaumer Ricaurte at the International Center for Tropical Agriculture in Cali, Colombia, and Kenneth J. Boote, Jose A Clavijo, Li Zhang and Tara Bongiovanni at the University of Florida. We thank Don McCarty and Harry Klee for their critical reading of the manuscript and their valuable suggestions. We also like to express our gratitude to the reviewers who provided very insightful suggestion that improved the quality of the manuscript.

## CONTRIBUTIONS BY THE AUTHORS

## Figure Legends

**Figure 1. Observed vs. predicted time-to-flowering plots (1:1) of the experimental recombinant inbred family at the five sites.** Predicted times were obtained with the CROPGRO-Bean model (**a**), the static (**b**) and the dynamic (**c**) statistical mixed-effects models. The R^2^_Adjusted_ value was used for comparative evaluation of the models to take into account sample size and the number of model parameters. Notice that the R^2^_Adjusted_ for (a) is the average of the values calculated for each RIL that had observations at the five MET sites.

**Figure 2 Differential equation describing the rate of development (RF) towards time-to-flowering as a function of genotype (g) and environmental (Tmax, Tmin, DL and Srad) factors at five experimental sites (s)**. This equation is represented by a linear mixed effects function where: DL_sgt_ = day length (hours) experienced by genotype *g*^*th*^ on day *t* at site *s*, DL_m_ = mean day length across all five sites (12.37 h); Srad_sgt_ = solar radiation (Srad, MJ m^-2^ d^-1^) experienced by genotype *g*^*th*^ on day *t* at site *s*, Srad_m_ = mean solar radiation across all five sites (18.218 MJ m^-2^ d^-1^); Tmin_sgt_ = minimum temperature (°C) experienced by genotype *g*^*th*^ on day *t* at site *s*, Tmin_m_ = mean minimum temperature across all five sites (16.128 °C); Tmax_sgt_ = maximum temperature (°C) experienced by genotype *g*^*th*^ on day *t* at ste *s*, Tmax_m_ = mean maximum temperature across all five sites (27.458 °C); TF1_j_ to TF12_j_ = QTL operators for the *j*^*th*^ allele (Calima alleles = +1 and Jamapa alleles = −1). For the rate equation: 0.02351 (d^-1^) is the mean effect of the environmental factors.

**Figure 3. Validation tests of the TF dynamic simulation model**. Observed vs. predicted plots (1:1) of two data sets not involved in generating of the model. (**a**) Parental genotypes that were grown along the RIF at the five MET sites. (**b**) A subset of 100 RILs, along with the parents, grown in Citra, FL, in 2016.

**Figure 4. Sensitivity analysis of the Time-to-Flowering module using the parental genotypes: Calima and Jamapa**. (**a**) Simulations of temperature effects on TF according to Equations (5) and (6) under two day-lengths. (**b**) Simulation of day length effects on TF with Tmax and Tmin values of 26 and 18 °C, respectively.

**Figure 5. Density plots of Time-to-Flowering estimate of the RI family at the five experimental sites**. Frequency distribution of the experimental RI family (Green; n = 188) and the synthetic RI family comprising all 4,096 possible allelic combinations of the 12 QTLs as simulated by the dynamic module. These plots indicate that multiple allelic combinations can attain the same phenotype providing a choice of convenience to plant breeding programs.

## Author contributions

C.E.V. and J.W.J. conceived of the project and wrote the manuscript;

S.A.G. and M.S.B. developed the mixed-effects models; M.S.B. and J.W.J. developed the R and FORTRAN scripts for the dynamic module, respectively; M.J.C. estimated the model parameters of the RI family, and C.E.V. carried out the QTL analysis of the model parameters.

## Data and materials availability

All the data and computer code are available in the supplementary materials. Seeds will be made available upon request.

