## Supplementary File for "Dynamic Gene-Based Ecophysiological Models to Predict Phenotype from Genotype and Environment Data"

**SUPPLEMENTARY INFORMATION**

**Supplementary Figures**

- **Supplementary Fig. 1 Temperature simulations generated with the dynamic gene-based time-to-first-flower bean model – implementation of Eqs. [2] and [3].**
- **Supplementary Fig. 2 Photoperiod simulations generated with the dynamic gene-based time-to-first-flower bean model – implementation of Eqs. [2] and [3].**
- **Supplementary Fig. 3 Photoperiod simulations of real and synthetic genotypes created through substitutions of QTL alleles.**

**Supplementary Tables**

- **Supplementary Table 1 Geographical and environmental characteristics of the five experimental sites.**
- **Supplementary Table 2 Genotype-specific parameter (GSP) values.**
- **Supplementary Table 3 Summary of QTL mapping results.**
- **Supplementary Table 4 Allele operators of the TF QTLs.**

**Supplementary Text**

- **Flowering Rate Model- R code**
- **Rate of Flowering Sub-Module – Rate of Progress toward First Flower.**
- **Driver Sub-Module.**

**Supplementary Data Files**

- **Weather_daily.csv**
- **RIL_R1sata_weatherDAPtoFF.csv**
- **AllQTLcombo.csv**

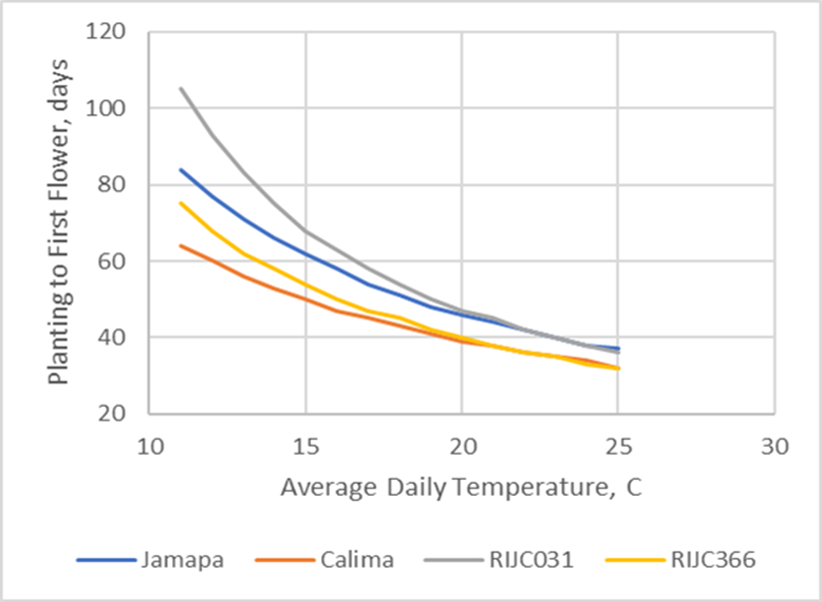

**Supplementary Fig. 1 Temperature simulations generated with the dynamic gene-based time-to-first-flower bean model – implementation of Eqs. [2] and [3].** The two parental genotypes Calima and Jamapa and two selected RILs shows different temperature dependencies. Notice the simulated transgressive behavior of RIJC031, which is in accordance with the observed data.

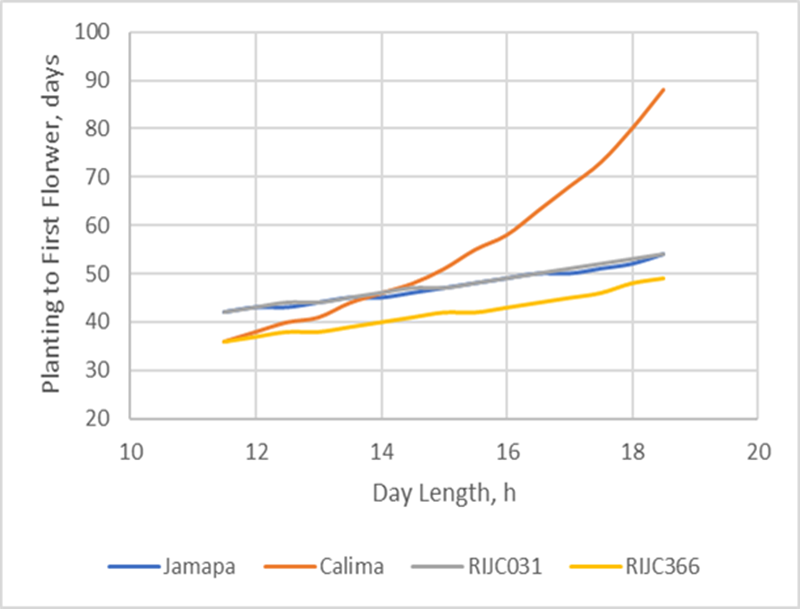

**Supplementary Fig. 2 Photoperiod simulations generated with the dynamic gene-based time-to-first-flower bean model – implementation of Eqs. [2] and [3.** The two parental genotypes Calima and Jamapa and two selected RILs display different photoperiod sensitivities Notice the simulated transgressive behavior of RIJC366, which parallels the observed data.

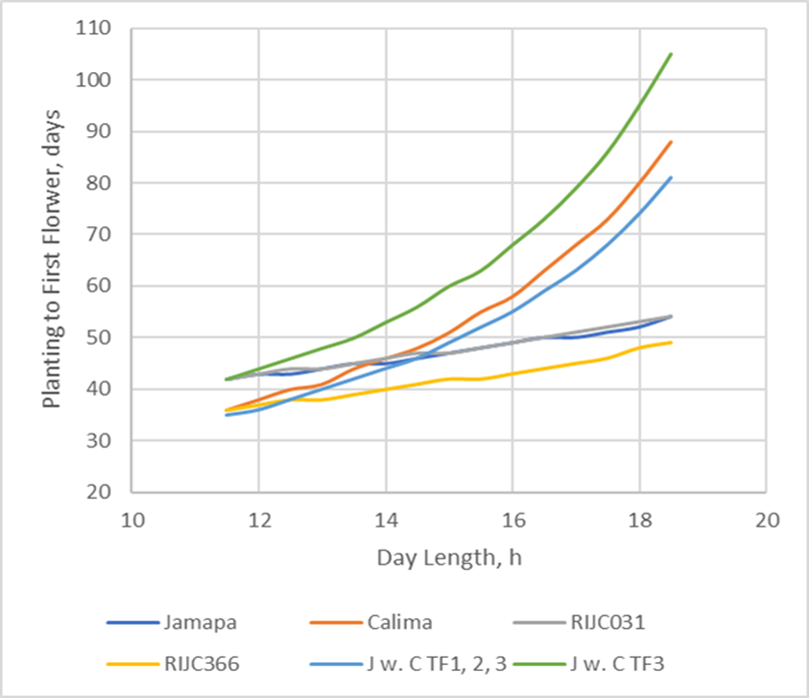

**Supplementary Fig. 3 Photoperiod simulations of real and synthetic genotypes created through substitutions of QTL alleles.** Results for the two parental genotypes Calima and Jamapa, two selected RILs and two synthetic lines. The response labeled “J w. CTF3” corresponds to a synthetic Jamapa genotype by substituting its TF3 allele with the Calima allele, which confers high sensitivity to long days. This single substitution conferred photoperiod sensitivity to Jamapa. Similar allelic replacement at two additional QTLs (TF1 and TF2), as shown by the response labeled “J w. C TF1,2,3”, increased the photoperiod sensitivity directly through TF3 and indirectly through interaction terms with the other QTLs.

| **Supplementary Table 1** **Geographical and environmental characteristics of the five experimental sites**. The recombinant inbred family was phenotyped for time-to-flowering at these sites. | | | | | | | | |
| --- | --- | --- | --- | --- | --- | --- | --- | --- |
| **Experimental Site** | **Site ID** | **MASL^a^** | **Latitude** | **Longitude** | **Day-Length**  **range (h)^b^** | **Solar Rad.^c^**  **MJ m^-2^ d^-1^** | **T_max_ (^o^C)^c^** | **T_min_ (°C)^c^** |
| Prosper, ND, USA | ND | 280 | 46° 53’ N | 97° 01’ W | 15:20 - 15:53 | 20.92 | 26.9 | 13.2 |
| Citra, FL, USA | CIT | 31 | 29° 24’ N | 82° 06’ W | 12:30 - 13:30 | 20.30 | 29.4 | 14.3 |
| Isabela, PR, USA | PR | 128 | 18° 28’ N | 61° 01’ W | 11:30 - 12:35 | 20.82 | 28.1 | 19.7 |
| Palmira, Colombia | PAL | 1000 | 03° 32’ N | 76° 18’ W | 11:56 - 11:58 | 14.17 | 28.2 | 19.6 |
| Popayan, Colombia | POP | 1800 | 02° 26’ N | 76° 36’ W | 12:08 - 12:11 | 15.14 | 24.6 | 14.0 |
| ^a^Meters Above Sea Level. ^b^During growing period. ^c^Average during the growing period. | | | | | | | | |

| **Supplementary Table 2 Genotype-specific parameter (GSP) values.** PL-EM (Planting to emergence), EM-FL (Emergence to flowering) and PPSEN (Photoperiod sensitivity) are GSPs associated with time-to-flowering in the DSSAT Bean-CROPGRO model. GSPs for the parents and the RIF were estimated via inverse modeling. | | | | | | | |
| --- | --- | --- | --- | --- | --- | --- | --- |
| **Genotypes** | **PL-EM** | **EM-FL** | **PPSEN** | **Genotypes** | **PL-EM** | **EM-FL** | **PPSEN** |
| CALIMA | 5.75 | 21.7 | 0.129 | RIJC0081 | 3.91 | 27.9 | 0.045 |
| JAMAPA | 4.57 | 30.3 | 0.027 | RIJC0082 | 4.79 | 27.3 | 0.039 |
| RIJC0003 | 4.79 | 21.1 | 0.131 | RIJC0129 | 4.79 | 27.7 | 0.036 |
| RIJC0004 | 5.76 | 28.6 | 0.053 | RIJC0130 | 4.79 | 27.2 | 0.019 |
| RIJC0005 | 4.73 | 23.7 | 0.135 | RIJC0131 | 3.92 | 23.9 | 0.080 |
| RIJC0006 | 3.97 | 21.6 | 0.111 | RIJC0135 | 4.81 | 31.4 | 0.025 |
| RIJC0007 | 6.86 | 27.6 | 0.080 | RIJC0136 | 3.90 | 28.8 | 0.043 |
| RIJC0008 | 4.79 | 22.1 | 0.162 | RIJC0137 | 3.90 | 23.2 | 0.125 |
| RIJC0009 | 3.97 | 29.6 | 0.043 | RIJC0138 | 3.90 | 28.4 | 0.041 |
| RIJC0011 | 6.76 | 32.4 | 0.071 | RIJC0140 | 4.88 | 23.0 | 0.147 |
| RIJC0012 | 4.73 | 22.8 | 0.122 | RIJC0141 | 5.73 | 28.2 | 0.050 |
| RIJC0013 | 4.74 | 22.9 | 0.098 | RIJC0142 | 3.94 | 33.7 | 0.058 |
| RIJC0014 | 5.78 | 22.6 | 0.142 | RIJC0144 | 5.88 | 29.6 | 0.068 |
| RIJC0015 | 3.90 | 22.9 | 0.100 | RIJC0145 | 4.79 | 29.5 | 0.041 |
| RIJC0016 | 4.60 | 31.5 | 0.042 | RIJC0146 | 4.73 | 27.7 | 0.050 |
| RIJC0017 | 4.79 | 28.8 | 0.112 | RIJC0147 | 5.74 | 31.0 | 0.068 |
| RIJC0019 | 5.79 | 25.7 | 0.088 | RIJC0148 | 4.79 | 25.9 | 0.124 |
| RIJC0020 | 4.79 | 23.6 | 0.060 | RIJC0149 | 5.77 | 28.2 | 0.098 |
| RIJC0021 | 4.60 | 23.9 | 0.051 | RIJC0150 | 5.82 | 23.9 | 0.081 |
| RIJC0022 | 5.75 | 31.0 | 0.087 | RIJC0151 | 5.86 | 27.8 | 0.060 |
| RIJC0024 | 6.10 | 28.7 | 0.161 | RIJC0201 | 4.69 | 24.0 | 0.056 |
| RIJC0025 | 5.79 | 26.9 | 0.103 | RIJC0202 | 5.91 | 31.3 | 0.117 |
| RIJC0027 | 4.60 | 27.0 | 0.104 | RIJC0203 | 4.79 | 27.6 | 0.057 |
| RIJC0029 | 4.79 | 22.0 | 0.095 | RIJC0204 | 5.83 | 28.1 | 0.051 |
| RIJC0032 | 6.26 | 27.9 | 0.075 | RIJC0205 | 4.79 | 26.1 | 0.051 |
| RIJC0045 | 5.74 | 24.8 | 0.137 | RIJC0206 | 3.90 | 27.8 | 0.059 |
| RIJC0046 | 4.99 | 26.9 | 0.057 | RIJC0207 | 3.90 | 27.9 | 0.075 |
| RIJC0047 | 4.81 | 22.2 | 0.141 | RIJC0208 | 3.90 | 33.3 | 0.071 |
| RIJC0048 | 4.79 | 29.1 | 0.057 | RIJC0209 | 5.82 | 29.0 | 0.061 |
| RIJC0059 | 4.79 | 28.1 | 0.104 | RIJC0210 | 4.79 | 28.1 | 0.086 |
| RIJC0061 | 4.79 | 30.1 | 0.082 | RIJC0212 | 5.43 | 25.6 | 0.115 |
| RIJC0065 | 4.79 | 26.9 | 0.064 | RIJC0213 | 5.43 | 22.1 | 0.153 |
| RIJC0066 | 5.75 | 23.5 | 0.155 | RIJC0214 | 5.73 | 24.0 | 0.120 |
| RIJC0067 | 4.79 | 26.6 | 0.066 | RIJC0216 | 6.49 | 26.8 | 0.047 |
| RIJC0069 | 3.90 | 28.5 | 0.039 | RIJC0217 | 3.90 | 22.6 | 0.013 |
| RIJC0070 | 4.60 | 27.8 | 0.062 | RIJC0219 | 6.78 | 27.9 | 0.094 |
| RIJC0071 | 3.97 | 25.8 | 0.161 | RIJC0220 | 3.91 | 28.2 | 0.049 |
| RIJC0072 | 4.73 | 29.7 | 0.046 | RIJC0223 | 3.91 | 24.8 | 0.150 |
| RIJC0073 | 3.90 | 28.0 | 0.084 | RIJC0224 | 3.90 | 26.7 | 0.033 |
| RIJC0074 | 3.90 | 23.5 | 0.122 | RIJC0225 | 5.77 | 24.7 | 0.074 |
| RIJC0076 | 4.69 | 28.1 | 0.039 | RIJC0228 | 5.77 | 24.3 | 0.050 |
| RIJC0078 | 3.90 | 22.0 | 0.053 | RIJC0229 | 3.92 | 28.1 | 0.043 |
| RIJC0079 | 5.76 | 27.1 | 0.053 | RIJC0230 | 4.58 | 28.9 | 0.033 |
| RIJC0080 | 3.91 | 21.7 | 0.076 | RIJC0231 | 5.91 | 24.4 | 0.088 |
| **Genotypes** | **PL-EM** | **EM-FL** | **PPSEN** | **Genotypes** | **PL-EM** | **EM-FL** | **PPSEN** |
| RIJC0232 | 4.79 | 33.1 | 0.088 | RIJC0320 | 3.90 | 30.1 | 0.097 |
| RIJC0233 | 4.79 | 23.9 | 0.100 | RIJC0322 | 4.85 | 23.0 | 0.107 |
| RIJC0234 | 6.38 | 24.3 | 0.125 | RIJC0324 | 5.67 | 29.1 | 0.035 |
| RIJC0235 | 4.99 | 30.5 | 0.058 | RIJC0326 | 5.75 | 25.4 | 0.048 |
| RIJC0236 | 5.66 | 28.1 | 0.066 | RIJC0327 | 4.79 | 22.8 | 0.130 |
| RIJC0237 | 4.79 | 27.4 | 0.059 | RIJC0328 | 6.49 | 28.3 | 0.060 |
| RIJC0238 | 3.90 | 27.6 | 0.040 | RIJC0330 | 5.43 | 25.6 | 0.056 |
| RIJC0239 | 5.35 | 23.7 | 0.091 | RIJC0333 | 5.74 | 21.9 | 0.128 |
| RIJC0243 | 4.79 | 28.3 | 0.069 | RIJC0334 | 4.79 | 31.3 | 0.063 |
| RIJC0245 | 4.78 | 22.5 | 0.101 | RIJC0335 | 4.79 | 22.1 | 0.093 |
| RIJC0246 | 5.86 | 24.3 | 0.099 | RIJC0337 | 3.90 | 28.4 | 0.041 |
| RIJC0247 | 3.90 | 27.5 | 0.088 | RIJC0339 | 5.76 | 31.7 | 0.045 |
| RIJC0248 | 4.79 | 23.0 | 0.179 | RIJC0340 | 6.38 | 25.6 | 0.116 |
| RIJC0249 | 4.79 | 29.7 | 0.052 | RIJC0342 | 5.83 | 27.0 | 0.039 |
| RIJC0250 | 4.18 | 23.1 | 0.137 | RIJC0343 | 7.36 | 26.3 | 0.059 |
| RIJC0251 | 4.80 | 23.4 | 0.056 | RIJC0344 | 4.03 | 30.4 | 0.061 |
| RIJC0252 | 5.75 | 24.1 | 0.149 | RIJC0346 | 3.91 | 28.0 | 0.084 |
| RIJC0253 | 5.87 | 24.7 | 0.106 | RIJC0347 | 4.73 | 34.5 | 0.055 |
| RIJC0254 | 3.91 | 23.1 | 0.118 | RIJC0348 | 4.75 | 24.1 | 0.139 |
| RIJC0255 | 3.91 | 26.0 | 0.105 | RIJC0349 | 3.92 | 30.3 | 0.073 |
| RIJC0257 | 4.57 | 21.4 | 0.094 | RIJC0350 | 4.79 | 28.7 | 0.054 |
| RIJC0259 | 5.75 | 28.3 | 0.084 | RIJC0352 | 4.03 | 27.5 | 0.041 |
| RIJC0260 | 5.34 | 23.9 | 0.127 | RIJC0353 | 4.60 | 27.7 | 0.050 |
| RIJC0261 | 5.76 | 26.9 | 0.123 | RIJC0355 | 4.79 | 27.4 | 0.082 |
| RIJC0262 | 5.76 | 32.6 | 0.067 | RIJC0356 | 3.90 | 28.0 | 0.068 |
| RIJC0264 | 4.14 | 29.7 | 0.087 | RIJC0357 | 4.93 | 31.7 | 0.028 |
| RIJC0301 | 3.91 | 28.7 | 0.094 | RIJC0358 | 5.72 | 31.8 | 0.100 |
| RIJC0302 | 5.75 | 25.9 | 0.081 | RIJC0360 | 4.79 | 29.2 | 0.097 |
| RIJC0303 | 4.62 | 30.1 | 0.042 | RIJC0362 | 4.73 | 29.1 | 0.050 |
| RIJC0306 | 3.90 | 26.9 | 0.072 | RIJC0363 | 4.79 | 23.3 | 0.145 |
| RIJC0311 | 4.79 | 23.8 | 0.134 | RIJC0364 | 5.74 | 21.3 | 0.114 |
| RIJC0313 | 3.90 | 24.8 | 0.104 | RIJC0366 | 3.91 | 24.3 | 0.020 |
| RIJC0314 | 4.79 | 26.5 | 0.131 | RIJC0367 | 3.90 | 23.2 | 0.081 |
| RIJC0316 | 3.91 | 24.0 | 0.045 | RIJC0368 | 4.79 | 23.2 | 0.105 |
| RIJC0317 | 5.68 | 29.4 | 0.059 | RIJC0370 | 4.79 | 30.1 | 0.035 |

| **Supplementary Table 3 Summary of QTL mapping results.** GSPs related to time-to-flowering in the DSSAT Bean-CROPGRO models were mapped in the RIF. The GSPs included PL-EM (Planting to emergence), EM-FL (Emergence to flowering) and PPSEN (Photoperiod sensitivity). | | | | | | |
| --- | --- | --- | --- | --- | --- | --- |
| **GSP** | **Chrom.** | **Marker Closest to QTL** | **Map Position (cM)** | **LOD** | **Effect %** | **QTL Overlap*** |
| PL-EM | 2 | DiM2-46 | 96.6 | 3.38 | 7.4 | - |
| PL-EM | 3 | DiM3-47 | 85.9 | 3.82 | 7.6 | - |
| PL-EM | 7 | DiM7-2 | 6.7 | 6.54 | 14.5 | - |
| EM-FL | 1 | DiM1-11 | 20.7 | 16.50 | 28.5 | TF1 |
| EM-FL | 1 | Fin | 41.6 | 26.40 | 45.0 | TF2 |
| EM-FL | 3 | DiM3-27 | 48.4 | 9.10 | 4.7 | TF6 |
| EM-FL | 11 | DiM11-9 | 18.3 | 3.42 | 2.2 | - |
| PPSEN | 1 | DiM1-29 | 59.8 | 23.31 | 53.2 | TF3 |

*Indicates whether the QTL for a GSP overlaps with a “time-to-flowering” QTL reported earlier by Bhakta et al.**^17^**

| **Supplementary Table 4 Allele operators of the TF QTLs.** These operators are used in the simulation of temperature and day length responses through the implementation of the fixed-effects model function described by Eq. [2]. The J-CTF3 and J-CTF1, 2, 3 correspond to Jamapa synthetic genotypes carrying Calima allele replacements at QTLs TF1 and TF1,2,3, respectively. | | | | | | | | | | | | |
| --- | --- | --- | --- | --- | --- | --- | --- | --- | --- | --- | --- | --- |
| **Genotype** | **TF1** | **TF2** | **TF3** | **TF4** | **TF5** | **TF6** | **TF7** | **TF8** | **TF9** | **TF10** | **TF11** | **TF12** |
| Calima | 1 | 1 | 1 | 1 | 1 | 1 | 1 | 1 | 1 | 1 | 1 | 1 |
| Jamapa | -1 | -1 | -1 | -1 | -1 | -1 | -1 | -1 | -1 | -1 | -1 | -1 |
| RIJC031 | -1 | -1 | -1 | -1 | 1 | 1 | 1 | -1 | -1 | 1 | 1 | 1 |
| RIJC366 | 1 | 1 | -1 | -1 | 1 | 1 | 1 | -1 | 1 | -1 | -1 | -1 |
| J-CTF3 | -1 | -1 | 1 | -1 | -1 | -1 | -1 | -1 | -1 | -1 | -1 | -1 |
| J-CTF1,2,3 | 1 | 1 | 1 | -1 | -1 | -1 | -1 | -1 | -1 | -1 | -1 | -1 |

```
Flowering Rate Model- R code

rm(list=ls())


## loading R libraries
library(lmerTest)
library(ggplot2)

### Setup your directory location
setwd("PathToYourDirectory")

##Read input data
we <- read.table('Weather_daily.csv',header=T)
r1 <- read.table('R1data_weatherDAFtoFF.csv',header=T)
TFcombo <- read.table('AllQTLcombo.csv',header=T)

## Part 1 - model parameter estimation

#mean for centering
sradMean <-mean(r1$Srad)
dayMean<- mean(r1$Day)

tminMean<- mean(r1$Tmin)
tmaxMean<- mean(r1$Tmax)


#centering continuous variable
r1$Srad_c<-r1$Srad-sradMean
r1$Day_c<-r1$Day-dayMean

r1$Tmin_c<-r1$Tmin-tminMean
r1$Tmax_c<-r1$Tmax-tmaxMean

r1$R1rate <- 1/r1$R1    ### 1/TFi variable creation

###### Linear Mixed-Effect Model for Flowering Rate 1/TF #############################################
modelr1rate = lmer((R1rate)~Tmax_c+Tmin_c+Day_c+Srad_c+
                     TF1+TF2+TF3+TF4+TF5+TF6+TF7+TF8+TF9+TF10+TF11+TF12+TF1*TF2+
                     Srad_c*TF12+Day_c*TF1+Tmax_c*TF5+Tmin_c*TF3+Day_c*TF3+
                     (1|RIL),data=r1)

summary(modelr1rate)
anova(modelr1rate)
##output fixed effect
#write.table( summary(modelr1rate)$coefficients,"fixedeffect.txt" , quote=F,row.names = F,sep="\t")

#output random effect
ranef(modelr1rate)


## predicting the R1 using Environment and QTL info from SAME DATASET used for model building and parameter estimation
r1$predRate <-predict(modelr1rate,newdata=r1)
summary(lm((r1$R1rate)~r1$predRate))$adj.r.squared
R.sqRateTf<-summary(lm((r1$R1rate)~r1$predRate))$adj.r.squared

## plotting observedRate vs predictated Rate
b<-ggplot(r1,aes(R1rate,predRate,color=SITE)) +
  scale_x_continuous(limits = c(0.005, 0.045)) +
  scale_y_continuous(limits = c(0.005, 0.045)) +
  geom_point(size=2.5)+theme_bw(base_size=16) +
  geom_abline(slope=1, intercept=0)+labs(x="Rate of Flowering",y="Predicted Rate of Flowering")
b<- b+ annotate("text", x = 0.015, y = 0.04, label = paste("Rsqare=",round(R.sqRateTf,2),sep=""))
b


## PART 2a

########################################################################################################
# Estimating daily flowering rate for each RIL at each SITE  or for ALL QTL combo - use line 71 or 72  START
######################################################################################################

r1 <- read.table('R1data_weatherDAFtoFF.csv',header=T)
#r1<-read.table("AllQTLcombo.csv",header=T)

predictionlist <- list()
predictionlist[[1]] =  rbind(predictionlist,c("SITE","RIL",'DayCount','CounterR1','dailyRateR1','stepper')) # first list row in prediction table--will be joined at the end of loop
#predictionlist[[1]] =  rbind(predictionlist,c("SITE","QTLCOMBO",'DayCount','CounterR1','dailyRateR1','stepper')) # first list row in prediction table--will be joined at the end of loop
rowcount = 1
rowcount = 1
stepper <-1
for (s in c("ND","CIT","PAL","POP","PR")){

  r2<-subset(r1,r1$SITE==s)  # subsetting RIL data file
  #r1<-subset(TFcombo,TFcombo$SITE==s)  ## subsetting QTL COMBINATION FILE
  we1 <- subset(we,SITE==s)  # subsetting weather data file

  ##looping over SITE,RIL,DAY
  for (i in 1:nrow(r2)) {
    SITE <- c(as.character(r2$SITE[i]))
    rsite <- SITE
    RIL <- c(as.character(r2$RIL[i]))
    o_R1 <- c(r2$R1[i])
    #QTLCombo <- c(r2$QTLCOMBO[i])
    TF1 <- c(r2$TF1[i])
    TF2 <- c(r2$TF2[i])
    TF3 <- c(r2$TF3[i])
    TF4<- c(r2$TF4[i])
    TF5<- c(r2$TF5[i])
    TF6 <- c(r2$TF6[i])
    TF7 <- c(r2$TF7[i])
    TF8 <- c(r2$TF8[i])
    TF9 <- c(r2$TF9[i])
    TF10 <- c(r2$TF10[i])
    TF11 <- c(r2$TF11[i])
    TF12 <- c(r2$TF12[i])

    #we1 <- subset(we,SITE==rsite)
    CounterR1 = 0
    DayCount = 0


    for (j in 1:nrow(we1)){
      DayCount <- we1$DAP[j]

      # Adjusting weather variables
      Srad_c <- c(we1$Srad[j])-sradMean
      Day_c <- c(we1$DAYLhr[j])-dayMean
      Tmax_c <- c(we1$Tmax[j])-tmaxMean
      Tmin_c <- c(we1$Tmin[j])-tminMean
      r1day <- as.data.frame(SITE,RIL,TF1,TF2,TF3,TF4,TF5,TF6,TF7,TF8,TF9,TF10,TF11,TF12,Srad_c,Tmin_c,Tmax_c,Day_c)

      ## Estimating daily flowering rate gain based on "modelr1rate" model
      flowerNow<-predict(modelr1rate,re.form=NA,newdata=r1day)  ## prediction with random effect as ZERO
      CounterR1 = CounterR1 + flowerNow[[1]]  ## counting cumulative flowering rate
      dailyRateR1 = flowerNow[[1]]
      rowcount = rowcount + 1
      predictionlist[[rowcount]] <- c(SITE,RIL,o_R1,DayCount,CounterR1,dailyRateR1,stepper)
      #predictionlist[[rowcount]] <- c(SITE,QTLCombo,DayCount,CounterR1,dailyRateR1,stepper)

      ## exiting loop when rate reaches 1.02
      if (CounterR1[[1]] >1.02){break}
    }

    stepper = stepper +1
    print(c(SITE,stepper))
  }
}

predictionRate <- as.matrix(do.call("rbind", predictionlist)) # converting list to matrix
colnames(predictionRate)<-predictionRate[1,]  # fixing header row names
predictionRate<-predictionRate[-1,] # removing old header row

##writing daily rate prediction for TF QTL allele combination - (large output file)
write.csv(predictionRate,"Daily_rate_prediction_outputRIJC.csv",row.names=F,quote = F)
#write.csv(predictionRate,"Daily_rate_prediction_outputALLQTLCombo.csv",row.names=F,quote = F)

#############################################################
# Estimating daily flowering rate for each RIL at each SITE  END
############################################################


###  Part 2b ###

###################################################################
# Estimating the day on which flowering rate is estimated to be 1.0 START
##################################################################

##Read input file -  daily rate prediction output.csv file
predictionRate <- read.csv("Daily_rate_prediction_outputRIJC.csv",header=T)

##interpolating data to estimate Day count when Rate hits 1.0  -- USE FOR RIL
GuesstheDay <- NULL
for (i in unique(predictionRate$RIL)){
  subdata <- subset(predictionRate,RIL ==i & CounterR1 >0.94)
  for (j in unique(subdata$SITE)){
    frame <- subset(subdata,SITE == j)
    o_R1<- frame$o_R1[[1]]
    SITE <- frame$SITE[[1]]
    RIL <- frame$RIL[[1]]
    Rate_R1_DayPrediction <- spline(frame$CounterR1,frame$DayCount,xmin=1,xmax=1,n=1,method="natural")$y
    GuesstheDay = rbind(GuesstheDay, data.frame(SITE,RIL,Rate_R1_DayPrediction))
  }
}
### writing output to directory -- estimated day when the predicted flowering rate hit 1.0
write.table(GuesstheDay,"r1_rate1.0_Prediction_center_RIL.txt",row.names=F,quote = F,sep='\t')

##interpolating data to estimate Day count when Rate hits 1.0  -- USE FOR ALL QTL COMBINATION
predictionRate <- read.csv("Daily_rate_prediction_outputALLQTLCombo.csv",header=T)

GuesstheDay <- NULL
for (i in unique(predictionRate$QTLCOMBO)){
  subdata <- subset(predictionRate,QTLCOMBO ==i & CounterR1 >0.94)
  for (j in unique(subdata$SITE)){
    frame <- subset(subdata,SITE == j)
    o_R1<- frame$o_R1[[1]]
    SITE <- frame$SITE[[1]]
    QTLCOMBO <- frame$QTLCOMBO[[1]]
    Rate_R1_DayPrediction <- spline(frame$CounterR1,frame$DayCount,xmin=1,xmax=1,n=1,method="natural")$y
    GuesstheDay = rbind(GuesstheDay, data.frame(SITE,QTLCOMBO,Rate_R1_DayPrediction))
  }
}
### writing output to directory -- estimated day when the predicted flowering rate hit 1.0
write.table(GuesstheDay,"r1_rate1.0_Prediction_center_ALLQTLCOMBO.txt",row.names=F,quote = F,sep='\t')

###################################################################
# Estimating the day on which flowering rate is estimated to be 1.0 END
##################################################################

#END
```

INPUT DATA FILES

‘Weather_daily.csv’

‘R1data_weatherDAPtoFF.csv’

‘AllQTLcombo.csv’

**Rate of Flowering Sub-Module.** This sub-module computes the daily rate of progress from planting to the day when the first flower appears on the plants. The FORTRAN source code for the RF Module is listed below. The “!” on lines indicate that the line is a comment (for example, see line 2 below). This module is run once per day to compute the rate of development for the specific genotype being simulated in the day’s environmental conditions. Referring to line numbers in this code, Line 18 contains the environmental inputs into this module for each day, which are day length (DL_i_), solar radiation (Srad_i_), and maximum and minimum temperatures (Tmax_i_ and Tmin_i_). Another key input to the module is the vector (TF), which has the genetic information (QTL allele operators) for the RIL being simulated. This Line 18 also shows the output that is communicated back to the main driver, SumRF_ij_, which is computed by accumulating the daily values of RFij for the RIL, starting at the time of planting when SumRF_ij_ = 0.00. Lines 26 – 30 set this initial value on the first day of the simulation, assuming that the program starts on the day of planting. Lines 36 - 40 set the mean values of environmental variables across all five locations as experienced during each RIL’s developmental period between emergence and first flower appearance.

The equation that computes daily rate of development for the genotype and environmental conditions being considered is given in Lines 45 – 67. The equation shown in the **RF Module** was based on Eqs. [2] and [3]. This new dynamic version describes more of the variability in the data than did the original mixed-effects linear model from Bhakta et al. (2017). Further, this dynamic version is critical to its implementation in other environments where future weather is not known in real-time predictions and where environmental conditions differ from those in the multi-environment trials used to estimate the model. Also, note that one can replace the model shown here with another function that computes each day's rate of progress toward first flower, such as using the original bean rate of progress toward flowering model**^13^** or the dynamic gene-based model published by Wallach et al.**^31^** Finally, Lines 85 – 86 in the **RF Module** compute SumRF_ij_, the summation of the daily development rates for environment *i* and genotype *j*, the integration step given in Eq. [3] in the paper. The remaining lines in this module (92 – 140) define variables to help document the program; these lines are not executed when the bean model is run.

**Fortran Code for the Rate of Flowering Sub-Module.** Below is a listing of the FORTRAN computer code for computing daily rates of development after emergence up to the time of first flower.

```
!   Written, 2017-01-05 by Jim Jones
!   With assistance from CH Porter
!   Based on linear mixed effects models developed by Mehul Bhakta (Equations [2] and [3].

!   Module for Rate of Development toward first flower in Common Bean, using G, E, and G x E inputs
!     Integrating RF vs. time to simulate first flower event
!   Computes daily rate of development, accumulates it and passes
!      the cumulative value back to the calling main program
!   Inputs to this module are daily day length (DLi, h),
!      daily maximum temperature (Tmaxi, C), daily minimum temperature (Tmini, C),
!      daily mean solar radiation (Sradi, MJ/m2), QTLs (TFi), and prior cumulative development (SumRFi)
!   Based on data and relationships from Vallejos NSF project, using data from 5 locations,

!   Used M. Bhakta RF daily model fitted to RF = (1/TF) depending on E, G, and GxE factors
!------------------------------------------------------------------------

       Subroutine RFlower_rate (DLi, Sradi, Tmaxi, Tmini, TF, SumRFij)

        Real, intent(in) :: DLi, Sradi, Tmaxi, Tmini !daily weather data
        Real, dimension(70) :: TF                    !Vector of cultivar parameters, genotype i
        Real, intent(out) :: SumRFij                 !progression towards anthesis, env i, genotype j
        Real RFij, RFmean, DLm, Sradm, Tmaxm, Tminm

        logical first
        data first /.true./
        if (first) then
          SumRFij = 0.0
          first = .false.
        endif

!Averages across 5 env in datasets used to estimate model
!        Mean values of environmental variables, estimated during fitting process for specific dataset used
!            Values from M. Bhakta based on integrated rate model. Received 8/19/2018
         RFmean = 0.02351
         DLm    = 12.86318
         Sradm  = 18.2719
         Tmaxm  = 27.4529
         Tminm  = 16.11819
!------------------------------------------------------------------------
!       The dynamic gene-based mixed effects linear model, Equation [2]
!------------------------------------------------------------------------
!
          RFij = RFmean
     &    + 5.8840e-04 * (Tmaxi-Tmaxm)
     &    + 5.2210e-04 * (Tmini - Tminm)
     &    - 1.5630e-03 * (DLi - DLm)
     &    - 8.2470e-05 * (Sradi - Sradm)
     &    + 9.2750e-04 * TF(1)
     &    + 1.2560e-03 * TF(2)
     &    - 6.7040e-04 * TF(3)
     &    + 2.6840e-04 * TF(4)
     &    - 3.5190e-05 * TF(5)
     &    + 5.6700e-04 * TF(6)
     &    - 4.1480e-04 * TF(7)
     &    - 2.2070e-04 * TF(8)
     &    - 4.6380e-04 * TF(9)
     &    - 2.7200e-04 * TF(10)
     &    + 3.4190e-04 * TF(11)
     &    - 1.6040e-04 * TF(12)
     &    + 2.8710e-04 * TF(1) * TF(2)
     &    - 5.1290e-05 * (Sradi - Sradm) * TF(12)
     &    - 1.3290e-04 * (DLi - DLm) * TF(1)
     &    + 9.9320e-05 * (Tmaxi-Tmaxm) * TF(5)
     &    - 9.9550e-05 * (Tmini - Tminm) * TF(3)
     &    - 6.7300e-04 * (DLi - DLm) * TF(3)

!     &    - 2.8300e-06 * (Tmini - Tminm) * TF(2)????
!     &    - 1.1560e-04 * (DLi - DLm) * TF(12)????

!
!     Note that one can replace the above mixed effects linear model with
!       any function that computes each day's rate of progress toward 1st flower
!       including use of the original bean rate of progress toward flowering model
!------------------------------------------------------------------------
!     Compute time integral of development to pass back as cumulative
!       progress toward development each day
!     In the equation for computing SumRFij, the time step is assumed
!       to be 1.0 d for this module (fixed)
!------------------------------------------------------------------------
!     Limit rate of development to positive values;
!       initial value=0.0. When SumRFij first reaches 1.00,
!       flowering will occur
      if (RFij < 1E-5) RFij = 0.0
      SumRFij = SumRFij + RFij*1.0

      Return
      End Subroutine RFlower_rate

!------------------------------------------------------------------------
!   Variable Definitions:
!------------------------------------------------------------------------
!
!   RFmean = 0.02342 is the mean rate toward flowering (1/day) across the 5 site-years in the bean dataset
!   DLi = day length during time from sowing to flowering observed for the ith genotype (hours),
!   DLm = mean day length across all five sites, all genotypes (12.37 hrs),
!   Sradi = average solar radiation from sowing to flowering observed by the ith environment (Srad, MJ/m2d),
!   Sradm = mean solar radiation across all sites, genotypes and transplanting dates in these data (18.218 MJ/m2d),
!   Tmaxi = daily maximum temperature each day during time from planting to first flower
!   Tmaxm = average of Tmaxi from transplanting to first flower across all genotypes, sites, years, (27.458 °C),
!   Tmini = daily minimum temperature each day during time from planting to first flower
!   Tminm = mean of the daily minimum temperatures means across all treatments, from planting to 1st flower (16.128)
!   TF(1):TF(70) = alleles at QTL TF1 :TF70 in jth genotype
!------------------------------------------------------------------------
!   INPUTS to Module:
!------------------------------------------------------------------------
!   E(Daily Environmental variables): DLi, Sradi, Tmaxi, Tmini
!   G (Genetic Variables): TF(1) thru TF(70); only 12 are
!       used in the linear model developed by Bhakta et al. for Bean
!   The numerical coefficients in the equation were developed for this
!     specific bi-parental population. These are either coded in the equation
!     above as constant numerical values
!     that cannot be changed by users for other genotypes
!     Changing them would cause results outside the confines of the data
!     used to estimate them. One can change TFj to compute TF for a particular line
!   Initial value of progress toward development
!     SumRFij0 (initially, SumRFij0 = 0.0 at time of transplanting)
!------------------------------------------------------------------------
!   OUTPUTS from Module, Dynamic Variables:
!------------------------------------------------------------------------
!     SumRFij = current progress toward first flowering from emergence
!      (dimensionless), integral of RFij from crop emergence or
!      planting to current day (environment i, genotype j)
!     The starting time for accumulation of time to first flower depends
!      on the data used to fit the model. Ideally, it will be emergence.
!------------------------------------------------------------------------
!   Local variables:
!------------------------------------------------------------------------
!     RFij = Daily rate toward development, fraction such that when integrated over time and a value of 1.0 is reached (or exceeded
!           First Flower occurs on that day, this is the mixed effects dependent variable
!     RFij = Daily rate of progress from planting to first flower appearance for selected
!           genotype (j) and environmental factors on the current environment & day (i)
```

!------------------------------------------------------------------------

**Driver Sub-Module.** This sub-module controls the sequence of calculations as it runs the *Rate of Flowering Sub-Module.* Thus, it implements the dynamic bean rate of flowering model and demonstrates how the model uses the MET data to predict time to first flower. In this implementation, the **Driver Sub-Module** was designed to conduct a simple sensitivity analysis to study specific genotypic responses to varying day lengths and daily temperatures. This module produces output files that have results of time to first flower for each specified environmental condition for a given genotype. One can use this module to identify the allelic combination(s) that will determine a specific time to first flower under predefined environmental conditions, including those predicted by climate warming models. These output files can be used to produce graphs of time to first flower responses as a function of temperature and/or day length conditions.

The module first sets the genetic values for the RIL to be simulated (Lines 33 – 44). It then opens a file into which model outputs are to be written and writes text headers for variables that will be written to the files (Lines 52 – 72). Lines 74 – 83 set initial conditions for a new combination of G and E. Note that one could also read daily values from a data file in order to simulate time to flower under time -varying weather conditions (Lines 46 – 51). Lines 88 - 100 is where the daily weather conditions are specified for each day of each intended combination of G and E for the simulation. Because this program is designed to simulate time to first flower over a range of environments, Lines 108 – 129 contain loops to set values of day length (Lines 108 – 112) and temperature (Lines 114 – 122). For each combination of day length and temperature, the model simulates time of flowering for the selected genotype. Line 138 calls the RF module to run the model. Line 143 determines when the SumRF_ij_ first exceeds 1.0 for a given set of G and E conditions, and this day is the day when the first flower occurs. Lines 145 – 152 write outputs for the simulated conditions. This program loops back through the driver module to simulate the selected combinations of day length and temperature. To simulate a different RIL, the QTLs for the specific genotype must be specified in Lines 33 – 44. There are 12 QTL allele operators to set, which a value of +1 for each QTL representing the Calima parent and a value of -1 for the Jamapa parent. Other RILs have different combinations of +1 and -1 numerical values.

**Driver Sub-Module Code.** The driver main flowering sub-module contains the FORTRAN code listing developed to simulate dynamic time to first flower model using daily weather value inputs of day length, maximum and minimum temperature, and solar radiation. Comment lines have an “!” at the front of the line or in front of the comment on a line.

```
!******************************************************************************
!*   Driver Module for running the RF Module to predict time of flowering     *
!*   Written, 2016-09-27 CH Porter, Jim Jones                                 *
!******************************************************************************
Program Main
!------------------------------------------------------------------------
  implicit none

  integer DOY, SDat, YR, Fdoy, RunNo
  integer Tmaxi_N, Tmini_N, DLi_N, k, m, i

  real DLi, Sradi, Tmaxi, Tmini, ADAP
  real DLi_incr, DLi_start
  real Tmaxi_incr, Tmaxi_start
  real Tmini_incr, Tmini_start
  real SumRFij
  real, dimension(70) :: TF           !Vector of genotype markers / parameters
  character(8) CultivarID
  character(12) Weather
  character(1) text
!------------------------------------------------------------------------
! Sensitivity tests using each marker to be +1 (Calima) or to be -1 (Jamapa)
!------------------------------------------------------------------------

  CultivarID = "C=T1,2,3"   !Jamapa values are all = -1, Calima = +1
  TF = 0.0
! TF1	TF2	 TF3   TF4	TF5   TF6   TF7   TF8   TF9  TF10  TF11  TF12
!  1     1    1     1    1     1     1     1     1    1     1     1	    !Calima
! -1    -1   -1    -1    1     1     1    -1    -1    1     1     1         !RIJC031
!  1     1   -1    -1    1     1     1    -1     1   -1    -1    -1         !RIJC366
! -1    -1   -1    -1   -1    -1    -1    -1    -1   -1    -1    -1	    !Jamapa

  TF(1) = 1.0
  TF(2) = 1.0
  TF(3) = 1.0
  TF(4) = -1.0
  TF(5) = -1.0
  TF(6) = -1.0
  TF(7) = -1.0
  TF(8) = -1.0
  TF(9) = -1.0
  TF(10)= -1.0
  TF(11)= -1.0
  TF(12)= -1.0

!------------------------------------------------------------------------
!  Weather = "CCPO1201.WTH"
! initialization, Open Files for output
! NOTE: Could read daily weather data file here and replace constant env variables
!   with daily-changing values
!------------------------------------------------------------------------
  open(40,File='Sens.Temp.OUT',status='REPLACE')

  write(40,'(A,A,17(/,2X,A,F8.3))') &
    "Marker values for genotype: ", CultivarID, &
    "TF(1)  =", TF(1) , &
    "TF(2)  =", TF(2) , &
    "TF(3)  =", TF(3),  &
    "TF(4)  =", TF(4),  &
    "TF(5)  =", TF(5),  &
    "TF(6)  =", TF(6),  &
    "TF(7)  =", TF(7),  &
    "TF(8)  =", TF(8),  &
    "TF(9)  =", TF(9),  &
    "TF(10) =", TF(10), &
    "TF(11) =", TF(11), &
    "TF(12) =", TF(12)

  write(40,'(/,a)') "  Run#   Cultivar     DAYL      Srad     Tmax       Tmin      SumRFi      ADAP      ADOY"
  write(*,'(/,a)') "  Run#   Cultivar     DAYL      Srad     Tmax       Tmin      SumRFi      ADAP      ADOY"

!------------------------------------------------------------------------
! Set sowing/start day of year for flowering model to start
!  Initialize progress toward flowering, SumDRi, & Day of First Flower, Fdoy
!------------------------------------------------------------------------
  SDat = 90     !sowing date
  ADAP = 0.0    !counter for number of days from sowing to anthesis
  SumRFij = 0.0
  Fdoy = 0
  RunNo = 0
  YR = 00
!------------------------------------------------------------------------
!Set constant values for weather inputs for sensitivity analysis
!------------------------------------------------------------------------
! doesn't iterate Sradi (uses mean values); currently, only Tmaxi, Tmini and DLi
  Sradi = 18.218

  Tmaxi_start = 14.0
  Tmaxi_incr  = 1.0
  Tmaxi_N     = 15

  Tmini_start  = 6.0
  Tmini_incr  = 1.0
  Tmini_N     = 15

  DLi_start = 11.0
  DLi_incr  = 0.5
  DLi_N     = 15

  open(30,File="Anthesis.OUT",status='REPLACE')
  write(30,'(a)') "adap  yr doy  Srad  Tmax  Tmin  SumRFij"
!  write(*,'(a)') "adap  yr doy  Srad  Tmax  Tmin  SumRFij"
!------------------------------------------------------------------------
!Daylength (DLi) variations loop, m counter, and set
!Temperature variations simulation loop, k counter
!------------------------------------------------------------------------
  DLi = DLi_start
  do m = 1, DLi_N  !Daylength loop
!   Increment daylength
    DLi   = DLi + DLi_incr

    Tmaxi  = Tmaxi_start
    Tmini  = Tmini_start
!------------------------------------------------------------------------

    do k = 1, Tmaxi_N  !Temperature loop
!     Increment temperature
      Tmaxi  = Tmaxi + Tmaxi_incr
      Tmini  = Tmini + Tmini_incr
!------------------------------------------------------------------------

!     Initialize accumulators
      SumRFij = 0.0
      ADAP   = 0.0
      RunNo = RunNo + 1
!------------------------------------------------------------------------

!   daily time loop, each combination of T, DL
    do i = 1,200
    DOY = i    !starting day of weather
    if (DOY > SDat-1.and.SumRFij < 1.0)Then  !Accumulate development after planting date (Sdat)
     ADAP = ADAP+1

!------------------------------------------------------------------------
!   call flowering module to calculate & integrate daily development rate over time
     call RFlower_rate (DLi, Sradi, Tmaxi, Tmini, TF, SumRFij)
!------------------------------------------------------------------------

     write(30,'(f5.1,1X,i2,1X,I3.3,3f6.2,f8.4)') ADAP, YR, DOY, Sradi, Tmaxi, Tmini, SumRFij

        if (SumRFij > 1.0 .and. Fdoy < 1) then   !First flower occurs
      Fdoy = DOY
      write(40,'(i6,2x,A8, 3f10.1,F10.2,2X,F10.5,F10.1,I10)') RunNo,CultivarID,DLi,Sradi,Tmaxi,Tmini,SumRFij,ADAP,Fdoy
      write(*,'(i6,2x,A8, 3f10.1,F10.2,2X,F10.5,F10.1,I10)') RunNo,CultivarID,DLi,Sradi,Tmaxi,Tmini,SumRFij,ADAP,Fdoy
      endif
        endif
    enddo       !End daily time loop
!------------------------------------------------------------------------

        ADAP = 0.0
        SumRFij = 0.0
        Fdoy = 0

    enddo  !end of Temperature loop
  enddo    !end of Daylength loop
!------------------------------------------------------------------------

  Stop
End Program Main
```
